# mitoSplitter: A mitochondrial variants-based method for efficient demultiplexing of pooled single-cell RNA-seq

**DOI:** 10.1101/2023.04.16.537058

**Authors:** Xinrui Lin, Yingwen Chen, Li Lin, Kun Yin, Rui Cheng, Xiaoyu Wang, Ye Guo, Zhaorun Wu, Yingkun Zhang, Jin Li, Chaoyong Yang, Jia Song

**Author notes:** Correspondence (J.L.) and (C.Y.), (J.S.). Equal contribution (Xinrui Lin, Yingwen Chen, Li Lin).

## Abstract

Single-cell RNA-seq (scRNA-seq) analysis of multiple samples separately can be costly and lead to batch effects. Exogenous barcodes or genome-wide RNA mutations can be used to demultiplex pooled scRNA-seq data, but they are experimentally or computationally challenging and limited in scope. Mitochondrial genomes are small but diverse, providing concise genotype information. We developed “mitoSplitter”, an algorithm that demultiplexes samples using mitochondrial RNA (mtRNA) variants, and demonstrated that mtRNA variants can be used to demultiplex large-scale scRNA-seq data. Using affordable computational resources, mitoSplitter can accurately analyze 10 samples and 60,000 cells in 6 hours. To avoid the batch effects from separated experiments, we applied mitoSplitter to analyze the responses of five non-small cell lung cancer (NSCLC) cell lines to BET chemical degradation in a multiplexed fashion. We found the synthetic lethality of *TOP2A* inhibition and BET chemical degradation in BET inhibitor-resistant cells. The result indicates that mitoSplitter can accelerate the application of scRNA-seq assays in biomedical research.

## Introduction

In recent decades, single-cell RNA-sequencing (scRNA-seq) assays have become widely recognized as useful for biomedical research(1, 2). While scRNA-seq assays can robustly identify single-cell clusters and genetic markers corresponding to cell types(3, 4), the variation in different rounds of experiments can introduce undesirable noise to the data set, also known as batch effects. Moreover, the high cost of regular scRNA-seq experiments has limited their application in analyzing a large number of samples.

To address these challenges, various technologies have been developed to demultiplex data from mixed samples in scRNA-seq experiments using chemical probes or genetic engineering-based exogenous barcodes(5-8). However, while the use of chemical probes can effectively demultiplex data based on labeling, it is limited by the need for universal antibodies suitable for various analytes, which also increases the cost and sample preparation time. On the other hand, genetic engineering-based exogenous barcodes do not require any antibodies, but their application is limited to cell culture systems or model organisms, making them unsuitable for acquiring scRNA-seq data from mixed clinical samples. Furthermore, the experimental expertise required for creating exogenous barcodes and properly transforming them into cells hinders the widespread adoption of these techniques.

It is encouraging to note that the demultiplexing of scRNA-seq data can also be achieved by analyzing the natural genomic variants with(9, 10) or without(11-13) the use of whole genome sequencing (WGS) or whole exome sequencing (WES) as references. However, these methods are computationally expensive and complex due to the detection and analysis of single-nucleotide polymorphisms (SNPs) of the whole genome as endogenous barcodes. Additionally, assigning demultiplexed cell groups to specific samples still requires data from either WGS or WES, which can significantly increase costs. Therefore, there is a need for a demultiplexing method that offers comparable accuracy but with significantly improved computational efficiency.

Similar to the SNPs found in the nuclear genome, the mitochondrial genome also contains many variants. It has been validated recently that mitochondrial genome variants can be used as endogenous barcodes for lineage analysis(14, 15). And our previous study has shown that mitochondrial RNA variants can be used for single-cell lineage tracing(16). Based on this, we hypothesized that it would be feasible to demultiplex scRNA-seq data from mixed samples using the SNPs found in the mitochondrial genome. In this study, we developed a computational algorithm called “mitoSplitter” to demultiplex mixed scRNA-seq samples by referencing the SNPs found in mitochondrial RNA.

The mitoSplitter algorithm uses bulk RNA-seq data from mitochondria or whole cells, rather than relying on WGS or WES data, as references. This approach significantly reduces the experimental cost of building the reference data set and the computational cost of identifying the endogenous barcodes. Using a label propagation algorithm, mitoSplitter can reliably demultiplex scRNA-seq data from human cell lines or clinical samples with high accuracy and computational efficiency. This allowed us to analyze 60,000 cells from 10 donors in a single experiment. Moreover, we utilized mitoSplitter to identify a specific synthetic lethality of Bromodomain and extra-terminal (BET) and *TOP2A* inhibition in the non-small cell lung cancer (NSCLC) cell line A549.

## Results

### Working principle of mitoSplitter

Compared to other chromosomes, the mitochondrial genome is much shorter, with only 16,569 base pairs, making a greater sequencing depth feasible compared to a different chromosome (**Supplementary Figure 1**). The shorter length of the mitochondrial genome significantly reduced the required computational resources. In addition, the number of mitochondria in a cell can vary greatly, with some cells containing more than 2,000 mitochondria, resulting in a rich diversity of mitochondrial genome sequences. All of these characteristics make the mitochondrial genome an ideal genotyping material. To estimate whether the mitochondrial genome variants within a population were diverse enough for genotyping, we analyzed a published RNA sequencing data from Gene Expression Omnibus with accession number GSE107011(17). We found that the average mitochondrial SNP profile diversities between individuals were approximately 1.5-1.7 times greater than those found in other chromosomes (**Figure 1A-B**, the average mitochondrial SNP profile variations between individuals were also calculated using the whole genome sequencing data from the 1,000 genomes project (Phase 3)(18) detailed in **Supplementary Figure 2**). Therefore, we hypothesized that it is indeed feasible to use mtRNA variants for genotyping based on bulk RNA-seq data of either mitochondria or whole cells. Furthermore, we contend that it is possible to demultiplex the scRNA-seq data from mixed samples (from different donors) based on mtRNA variants.

**Figure 1.**
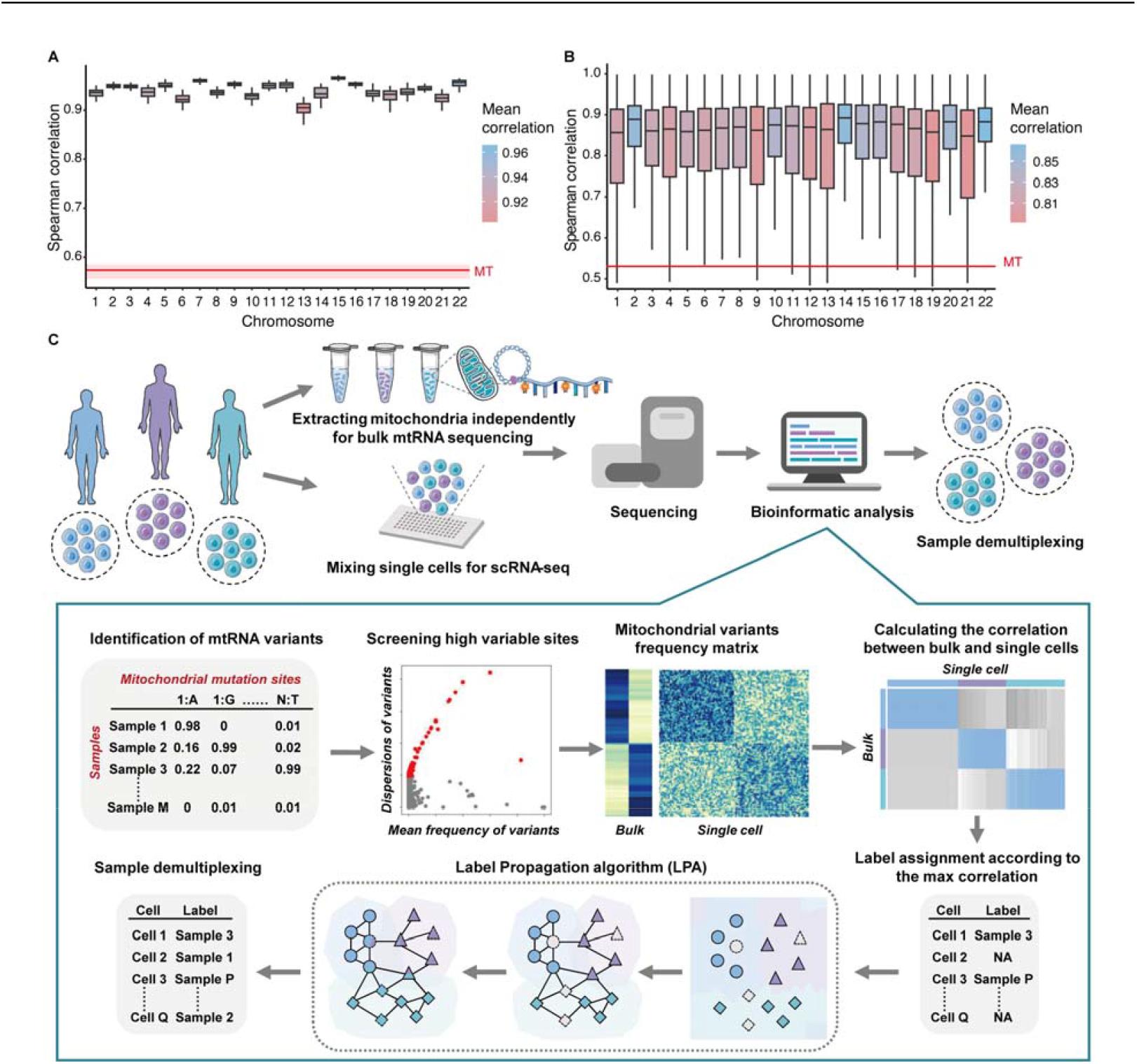
Mitochondrial variants as the optimal material for genotyping. (A) The mean Spearman correlation between all pairs of two individuals in a published RNA sequencing data based on mitochondrial or autosomal variant profiles. The red line represents the mean correlation of the mitochondrial variants between individuals, while the upper and lower pink boundaries represent the first and third quantiles. MT represents “mitochondrion”. These results demonstrated that mitochondrial variant profiles are significantly more variable between individuals than those of other chromosomes. (B) The mean Spearman correlation between pairwise individuals from a published RNA sequencing data based on variant profiles of mitochondrial or autosomal segments. An autosomal segment is composed of 13 adjacently located genes (equal to the gene number of the mitochondrial genome), and 100 segments were randomly extracted from each autosomal segment. The red line represents the mean correlation of the mitochondrial genome between individuals. MT represents “mitochondrion”. These results indicate that profiles of mitochondrial variants are significantly more variable than those of other chromosome segments containing the same number of genes. (C) Overview of mitoSplitter framework. SNPs were identified by aligning RNA-seq data to the reference genome for each sample. High-variability mitochondrial SNPs were identified based on variation frequency and dispersions in bulk RNA-seq, and used to genotype samples. The mitochondrial genotyping correlation between bulk samples and cells was used to determine the sample label of some single cells. Only cells with high bulk sample correction were labeled. Then, the label propagation algorithm (LPA) was employed to give sample labels to unlabeled cells based on single-cell mtRNA SNPs.

Subsequently, we established an algorithm named “mitoSplitter” to demonstrate these hypotheses (**Figure 1C**). Briefly, the mitochondrial SNPs were identified through alignment and analysis of the RNA-seq data (including both bulk and single-cell) to the reference genome for each sample. The highly-variable mitochondrial SNPs from the bulk data were selected for mitochondrial genotyping according to the variant’s specific frequency and dispersions. Once mitochondrial genotyping of samples was confirmed, a label representing which sample the individual cell was derived from was inferred according to correlation throughout bulk samples and cells, which was then included in the metadata of the scRNA-seq dataset. Due to the low depth of single-cell sequencing, some cells did not receive reads mapped to their selected highly variable sites (mitochondrial genotyping sites) and therefore did not receive labels. To overcome this challenge, a semi-supervised machine learning strategy, known as a label propagation algorithm (LPA), was integrated to transfer sample labels among cells according to specific mtRNA SNPs detected in single cells. Finally, the distribution of cells from different samples was visualized through the assignment of different colors to the samples on the TSNE plots. The source code and documentation for this computational algorithm, mitoSplitter, are available on Github (https://github.com/lnscan/mitoSplitter).

### Validation of mitoSplitter using virtually multiplexed scRNA-seq data

We first tested the reliability and robustness of mitoSplitter using two simulation datasets with different mixing scenarios involving 4 and 3 PBMC samples (detailed in **Supplementary Table 1 and 2**). The genotypes were obtained from the pooled scRNA-seq data for each donor (the sequencing depth of each donor’s mitochondrial genome is shown in **Supplementary Figure 3A and B**). We compared the accuracy, running time, and required memory of mitoSplitter to four published computational algorithms, including Demuxlet(9), Souporcell(13), scSplit(11), and Vireo(12). The results revealed that mitoSplitter demonstrated comparable or even higher accuracy (measured as AUC, see the details in **Supplementary Note 1**) than other tools for both datasets. However, the computation time and memory requirement of mitoSplitter were much lower than those of other methods (**Figure 2A**). Moreover, the accuracy of mitoSplitter’s demultiplexing was also validated using a TSNE plot based on mtRNA variants, where the “True” label from the original datasets and the “Predicted” label from mitoSplitter showed strong agreement (**Figure 2B**, the annotated cell types in the demultiplexed data are shown in **Supplementary Figure 4A and B**).

**Figure 2.**
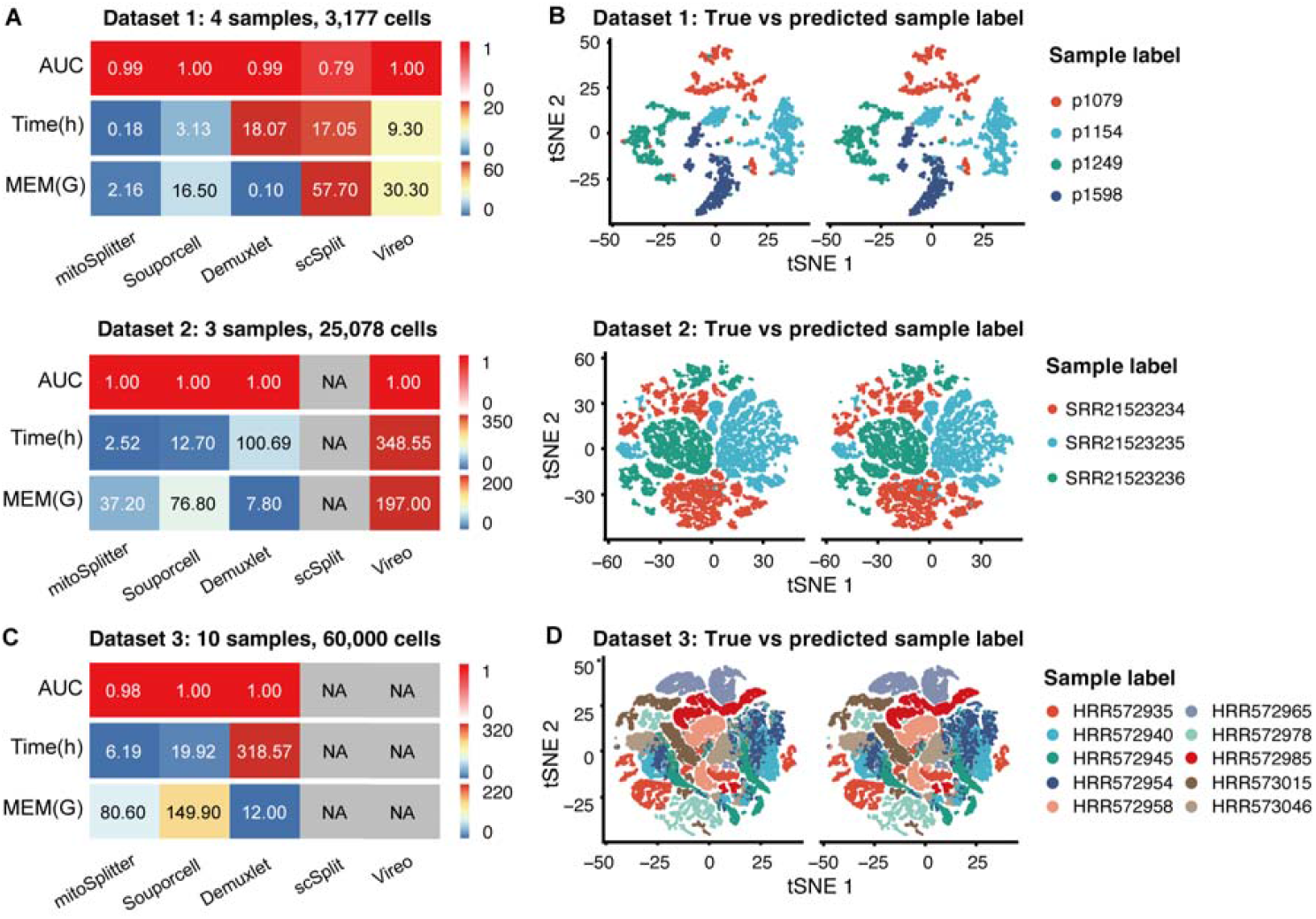
Performance of mitoSplitter on virtually multiplexed scRNA-seq data. (A) Comparison of the performance of mitoSplitter and other demultiplexing tools on two datasets with 4 and 3 samples, displaying AUC (area under the receiver operating characteristic curve), running time, and memory requirements for each tool. “NA” indicates that the tool was unable to complete the computation with the maximal memory (600G) on the server. (B) TSNE plot based on highly variable variants identified by mitoSplitter in two datasets with 4 and 3 samples. The left subplots were colored according to the true sample labels, while the right subplots were colored according to the sample labels predicted by mitoSplitter. (C) Comparison of the performance of mitoSplitter and other demultiplexing tools in a large dataset with 10 samples, displaying the AUC, running time, and memory requirements for each tool. “NA” indicates that the tool was unable to complete the computation with the maximal memory (600G) on the server. (D) TSNE plot based on highly variable variants identified by mitoSplitter in the large dataset. The left subplot was colored according to the true sample labels, while the right subplot was colored according to the mitoSplitter-predicted sample labels. “NA” indicates that the tool was unable to complete the computation with the maximal memory (600G) on the server.

We tested mitoSplitter and the other methods using a large virtually mixed scRNA-seq dataset comprising 60,000 cells from across 10 samples (**Supplementary Table 3**). MitoSplitter demonstrated decent accuracy within 6 hours on a server with ∼80 Gigabytes (G) of memory, making it an ideal tool for splitting large datasets. In contrast, scSplit and Vireo indeed failed to analyze this large dataset with maximal memory (600G) on our server (**Figure 2C**). The accuracy of mitoSplitter was further demonstrated in mtRNA variant-based TSNE plots (**Figure 2D**, the annotated cell types in the demultiplexed data are shown in **Supplementary Figure 4C**). These experiments on simulated datasets suggest that mitoSplitter may perform label-free scRNA-seq data demultiplexing with minimal computational resources.

### Validation of mitoSplitter using pseudo-mixed PBMC samples

We evaluated the reliability and robustness of mitoSplitter using newly generated scRNA-seq data and paired mitochondrial bulk RNA-seq data from the PBMC sample. Mitochondrial genotypes were identified from low-depth bulk RNA-seq data of mitochondria (∼3.7G per sample). PBMC samples from across 11 donors were analyzed using the Well-Paired-Seq assay(19) separately (**Supplementary Table 4**). The data from between 2 and 10 donors were virtually multiplexed by random selection of donors. Ten datasets were randomly created for each condition. The mitoSplitter algorithm was able to achieve an average AUC higher than 0.99 for different sample sizes and combinations (**Figure 3A, Supplementary Table 5**).

**Figure 3.**
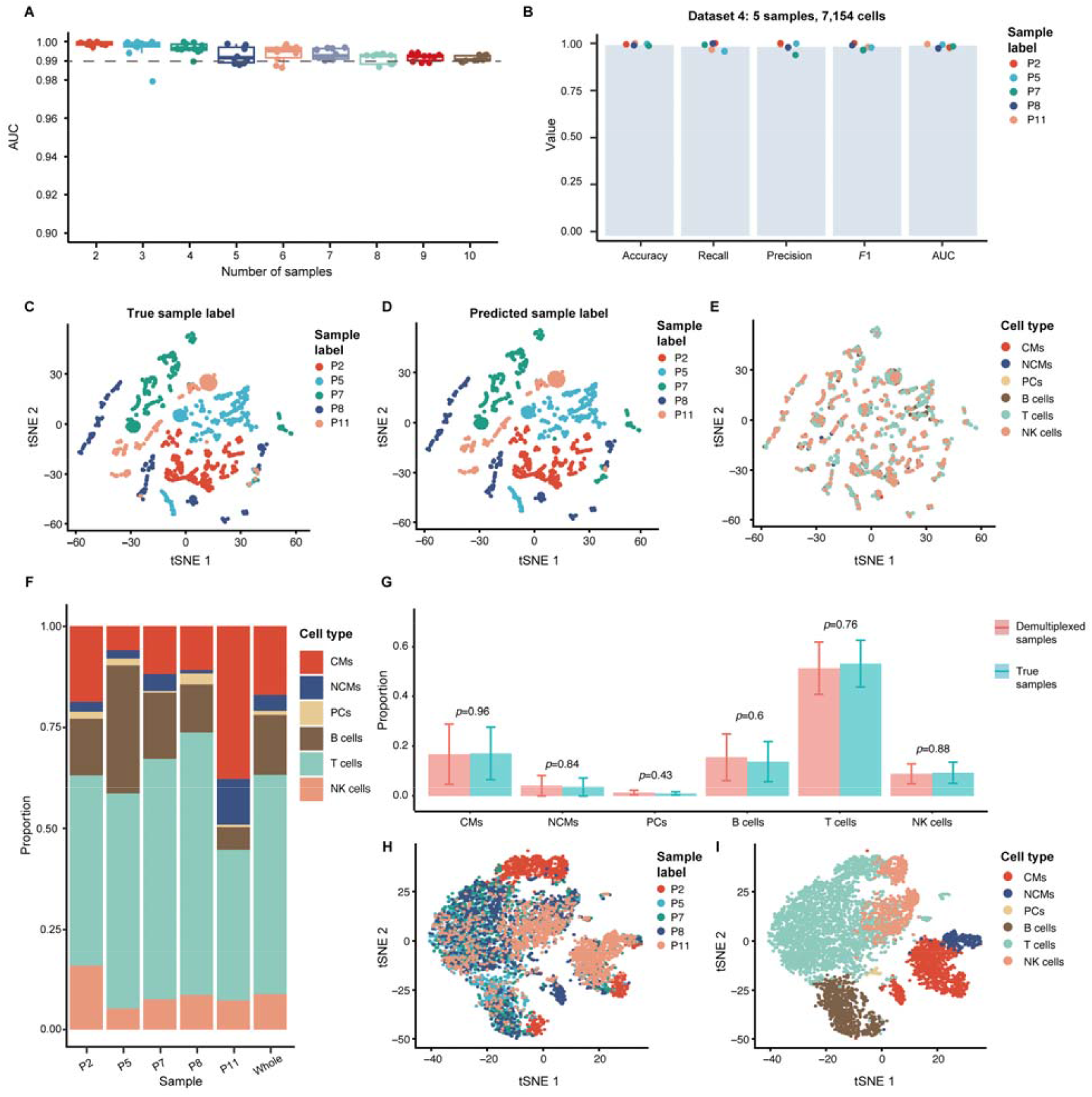
Performance of mitoSplitter on pseudo-mixed PBMC samples. (A) AUC values of mitoSplitter demultiplexing outcomes for varying numbers of mixed samples. (B) A comprehensive evaluation of mitoSplitter demultiplexing results for a dataset of five mixed samples, including demultiplexing accuracy, recall rate, precision, *F*1 score, and AUC for each sample. (C-E) TSNE plot generated using highly variable variants identified by mitoSplitter and colored based on true sample labels, predicted sample labels, and cell types, respectively. (F) Fraction of each cell type in the demultiplexed samples. (G) Comparison of cell type composition between demultiplexed and true samples using a paired t-test. (H-I) Gene expression-based TSNE plots colored based on predicted sample labels and cell types, respectively. Cell types include classical monocytes (CMs), non-classical monocytes (NCMs), plasma cells (PCs), B cells, T cells, and natural killer cells (NK cells).

We randomly selected a 5-donor pseudo-multiplexed dataset to demonstrate the performance details of mitoSplitter. The analysis of mitoSplitter on all 5 donors showed excellent accuracies (**Figure 3B**). To determine if the same biological conclusion can be drawn from the computationally demultiplexed data and experimentally separated data, we compared the downstream analysis results with the “True” and “Predicted” labels. The mtRNA variants-based TSNE plot illustrates the overall agreement of the data with the “True” and “Predicted” labels from mitoSplitter (**Figure 3C-D**). Moreover, the TSNE plots showed the universal distribution of 6 cell types from the 5 donors (**Figure 3E-F**). There was no significant difference in the proportion of each cell type with the “True” and “Predicted” labels in the data from the five donors, as indicated by a paired t-test (**Figure 3G, Supplementary Table 6**). Importantly, gene expression-based TSNE plots revealed remarkable batch effects, emphasizing the importance of performing scRNA-seq analysis with mixed samples (**Figure 3H-I**). These results underscore the ability of mitoSplitter to perform genotyping based on low-depth bulk RNA-seq data of mitochondria and demultiplex scRNA-seq data from up to 10 donors.

### Validation of mitoSplitter using experimentally multiplexed scRNA-seq data

Using exogenous barcodes as the gold standard, we verified the demultiplexing capability of mitoSplitter using multiplexed scRNA-seq data from antibody- or lipid-based multiplexing methods (**Figure 4A, Supplementary Table 7**). Notably, mitoSplitter achieved an AUC of 98%-100% from these data (**Figure 4B**), with a maximum time and memory consumption of 1.2 hours and 13.3G (**Figure 4C**), respectively. This demonstrates the robustness and high efficiency of this algorithm on experimentally mixed samples.

**Figure 4.**
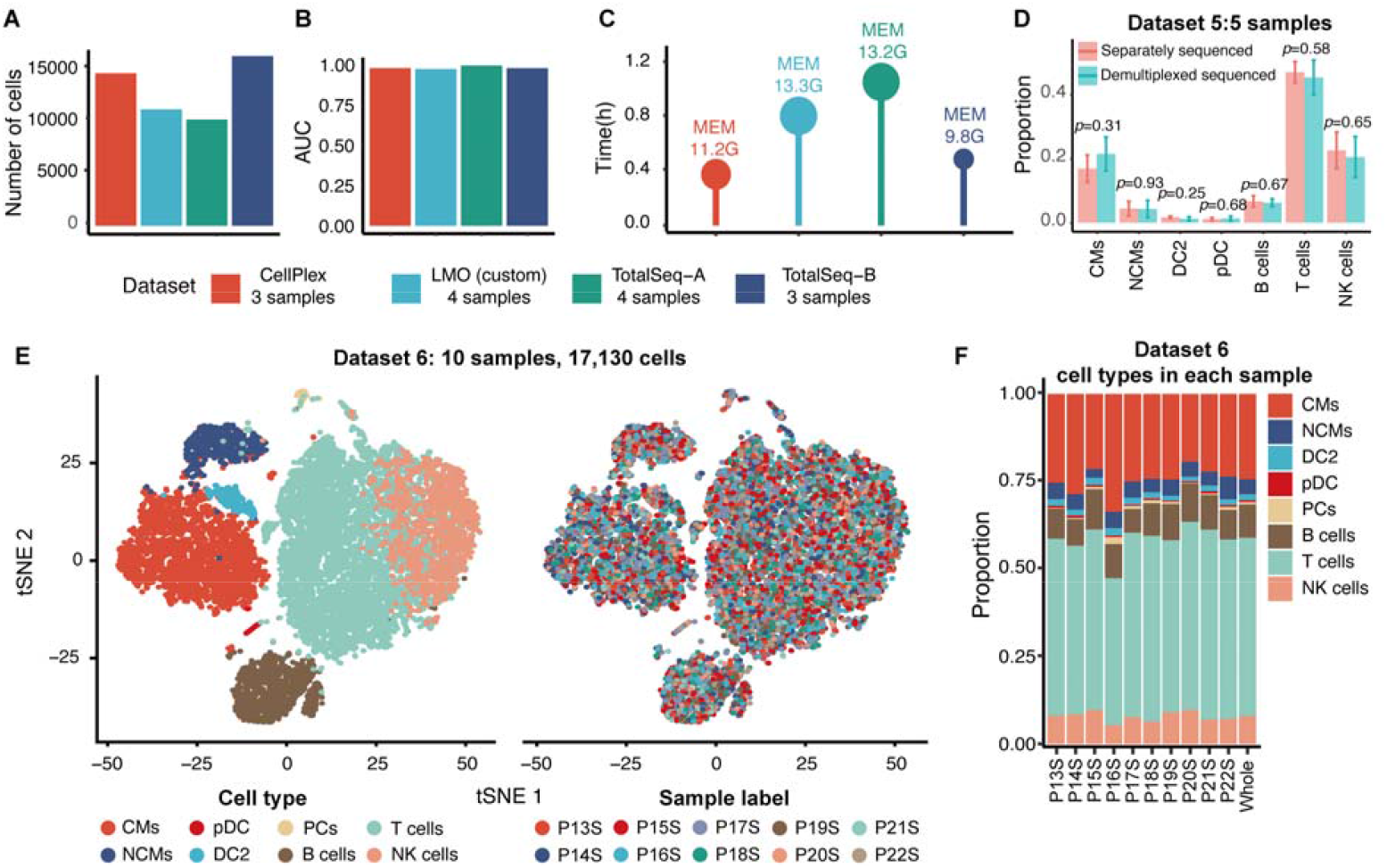
Performance of mitoSplitter on experimentally multiplexed scRNA-seq data. (A) The number of cells in each dataset. (B) The AUC of mitoSplitter on four multiplexed datasets with exogenous barcodes. (C) The running time and memory requirement of mitoSplitter on the same four multiplexed datasets with exogenous barcodes. (D) A paired t-test was used to compare the cell type composition of demultiplexed and separately sequenced samples. (E) The gene expression-based TSNE plot for multiplexed data with 10 samples. The left side is colored by cell types, while the right side is colored by predicted sample labels. (F) The proportion of each cell type in the demultiplexed samples. The cell types include classical monocytes (CMs), non-classical monocytes (NCMs), plasmacytoid dendritic cells (pDC), type 2 dendritic cells (DC2), plasma cells (PCs), B cells, T cells, and natural killer cells (NK cells).

To further determine the robustness of mitoSplitter using primary samples, 25,275 and 21,815 PBMCs from across 5 donors were analyzed in both separated and pooled fashion in parallel (**Supplementary Table 8**). MitoSplitter completed analysis in under 1.5 hours using 21.7G of RAM. 3,788 ± 738 cells were demultiplexed per sample (the TSNE results are shown in **Supplementary Figure 5A**). There was also no significant difference in the proportion of each cell type in the separated sequenced and demultiplexed samples from the five donors as verified by a paired t-test (**Figure 4D**, the cell type compositions are shown in **Supplementary Figure 5B and C**). These results suggest that the demultiplexed results from mitoSplitter were comparable to those of separated analysis.

Subsequently, we tested a large label-free scRNA-seq dataset consisting of 17,130 PBMCs from ten donors (**Supplementary Table 9**). MitoSplitter was able to complete the analysis in under 0.5 hour while using a negligible amount of RAM (35.6G). 1,443 ± 575 cells were detected per sample. We observed low batch effects between different donors in the gene expression-based TSNE plot (**Figure 4E**). The proportions of each cell type in the demultiplexed data from the ten donors were consistent with the existing knowledge of PBMC (**Figure 4F**). These findings indicate that mitoSplitter can reduce batch effects through a combination of as many as ten samples into a single scRNA-seq.

### scRNA-seq analysis of NSCLC using mitoSplitter identified synthetic lethality of a BET degrader and *TOP2A* inhibitor

Non-small cell lung cancer (NSCLC) is a significant cancer type worldwide. Previous research has demonstrated that the inhibition or degradation of bromodomain and extra terminal (BET) proteins may impede the proliferation of NSCLC cells(20-22). However, it has also been reported multiple times that the sensitivity of NSCLC cells to chemical inhibition or degradation of BET is highly variable(23-26). Consequently, the drugs which can induce the synthetic lethality to overcome the resistance to BET inhibitors are still missing. The potential reasons for the inconsistencies among these studies include the lack of analysis in single-cell resolution as well as the batch effects of different studies. Therefore, we were aiming to understand the mechanism for NSCLC’s resistance to BET inhibitor by employing multiplexed single-cell RNA-seq and mitoSplitter.

Firstly, we observed different effects of chemical degradation of BET through the treatment of ARV-771(27) on the proliferation rate of NSCLC cell lines (**Supplementary Figure 6A**). Interestingly, the A549 cell line exhibited significant resistance to ARV-771, prompting us to investigate the underlying resistance mechanism using mitoSplitter.

To mitigate batch effects, we performed multiplexed scRNA-seq analysis of five NSCLC cell lines, treated or untreated with ARV-771 for 48 hours, in a single experiment. The analysis revealed multiple clusters purely based on expression profiles (**Figure 5A**). Using bulk RNA-seq data of mitochondria, we assigned the cells to their origins and presented them in the TSNE plots (**Figure 5B, Supplementary Table 10**). Notably, ARV-771 treatment resulted in a decrease in the proportion of cells, except for A549 cells (**Figure 5C**).

**Figure 5.**
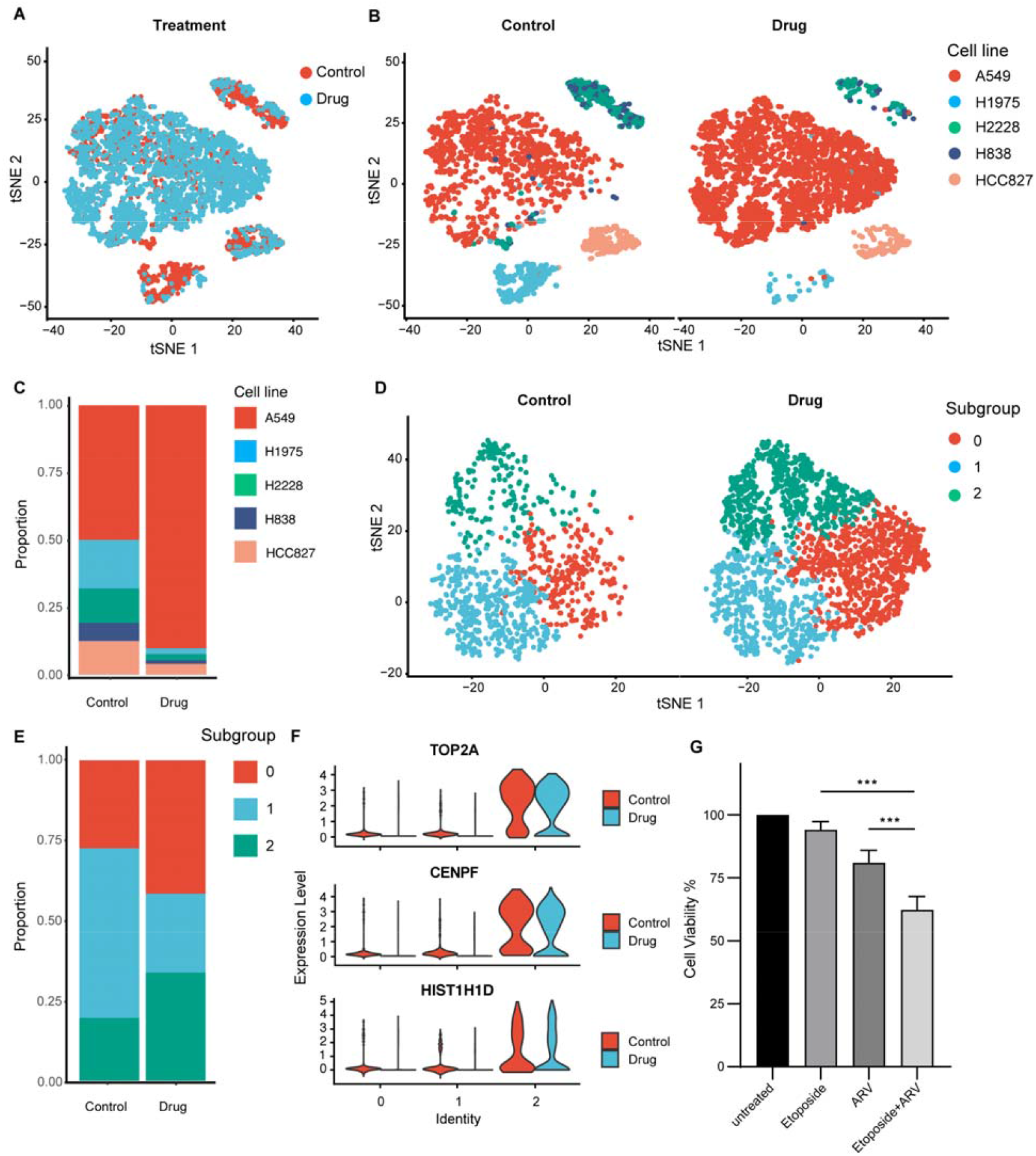
scRNA-seq analysis of NSCLC using mitoSplitter identified synthetic lethality of a BET degrader and *TOP2A* inhibitor. (A) The NSCLC cells are presented in a combined gene expression-based TSNE plot, colored by drug treatment status. (B) Two separate TSNE plots of NSCLC cells based on gene expression, colored by cell lines. The left plot shows cells that were not treated with the drug, while the right plot shows cells that were treated. (C) The proportion of cell lines in the sequencing results before and after drug treatment. (D) Two gene expression-based TSNE plots of A549 cells, where the left plot shows cells that were not treated with the drug, while the right plot shows cells that were treated. Cells are colored by subgroup types. (E) The proportion of subgroups in the A549 cell line before and after drug treatment, where the proportion of cells belonging to subgroup 0 and 2 increased after drug administration. (F) The expression of the top three down-regulated genes identified in A549 subgroup 1 across all subgroups. (G) Combined administration of Etoposide and ARV can significantly reduce the viability of A549 cells.

We investigated the mechanism of resistance to ARV-771 by performing a computational analysis of the scRNA-seq data from A549 cells. We observed the proliferation of two clusters (subgroup 0 and 2) in the presence of ARV-771 (**Figure 5D and E**) and identified a list of markers for these clusters (subgroup 1 compared to subgroups 0 and 2, **Supplementary Table 11**). Functional enrichment analysis of these markers revealed their association with mitosis and cell cycle (**Supplementary Figure 6B and C**). Notably, we observed an upregulation of *TOP2A* which is a well-known therapeutic target for NSCLC (**Figure 5F**). Importantly, the combined chemical inhibition of *TOP2A* with etoposide and ARV-771 synergistically inhibited the proliferation of A549 cells specifically (**Figure 5G, Supplementary Figure 6D**).

## Discussion

In the current study, we present a method called “mitoSplitter” for analyzing scRNA-seq data from multiple samples in a single experiment. This approach uses variants of mtRNA to deconvolute individual samples from pooled scRNA-seq data with reasonable computational time and memory usage. Our application of mitoSplitter to NSCLC revealed a synthetic lethality between *TOP2A* inhibition and BET chemical degradation.

The high cost of scRNA-seq limits its application in studying complex diseases, which often require analysis of a large number of samples. To increase throughput, one strategy is to multiplex multiple samples in a single experiment(28-31). However, existing strategies using endogenous or exogenous barcodes incur significant experimental or computational costs. In contrast, mitoSplitter only requires additional low-depth sequencing of mtRNA of each sample, which dramatically reduces the overall cost of multiplexed scRNA-seq assays. We anticipate that mitoSplitter will be useful for epidemiology and population genetics studies.

The batch effects among different experiments represent a major technical challenge for scRNA-seq analysis. Despite several efforts from the community(32-36), performing multiplexed scRNA-seq analysis in a single experiment is still the ideal method to overcome this issue. The existing methods of demultiplexing scRNA-seq data using either exogenous or endogenous barcodes require extensive experimental or computational efforts. By launching mitoSplitter, we are now able to demultiplex label-free scRNA-seq data with minimal computational expertise. Using mitoSplitter, biologists with limited experimental and computational capability can analyze mixed samples from scRNA-seq assays.

Before developing mitoSplitter, we were concerned that the lack of diversity in SNPs within the mitochondrial genome may hinder the demultiplexing of complicated samples. However, our study validated the robustness of mitoSplitter using various datasets containing 2-10 samples. Compared to other methods, mitoSplitter exhibited comparable or even higher accuracy in assigning cells to samples, while requiring less RAM and running time for all tested datasets. Notably, mitoSplitter performed similarly well on scRNA-seq data from both cell lines and primary cells.

The BET inhibitor or chemical degrader has shown potential as an anti-cancer agent for a wide range of cancer cells. Therefore, it is crucial to understand the mechanism of resistance to ARV-771 in A549 cells. Single cell RNA-seq is a useful assay for resistance studies since it can identify the features of resistant clones. But due to the batch effects of single cell RNA-seq assay, it is challenging to analyze the data of resistant and sensitive cells generated from separated experiments. In the current study, application of mitoSplitter on multiplexed single cell RNA-seq data revealed a clear clonal evolution event of the resistant A549 cells, when they were mixed other four sensitive cell lines and treated with BET degrader ARV-771. These results led us to further identify a previously unknown synthetic lethality between BET degraders and *TOP2A* inhibitors. However, the detailed mechanism underlying the emergence of the resistant clone requires further investigation.

Currently, mitoSplitter is limited by the requirement of mitochondrial genome information in the scRNA-seq data, making it unsuitable for single nucleus RNA-seq (snRNA-seq) analysis. As snRNA-seq is becoming a popular assay for analyzing frozen clinical samples, methods specifically designed for demultiplexing snRNA-seq data will need to be established in the future.

In conclusion, we developed the mitoSplitter assay to reliably and efficiently analyze multiple single-cell samples in one scRNA-seq experiment. With mitoSplitter, we have overcome many technical obstacles for biologists to perform scRNA-seq experiments with mixed samples. The availability of mitoSplitter will further accelerate the application of scRNA-seq assay in biomedical research.

## Material and Methods

### Cell preparation

Whole blood samples were obtained from donors with approval from the ethical committee of Xiamen University (Protocol No. KYX-2018-006). To isolate PBMCs, 3 mL of whole blood was fully mixed with 3 mL of 1×PBS and subsequently added onto 6 mL Ficoll-Paque Plus (GE Healthcare, 17-1440-02). After centrifugation at 2000 rpm at 15°C for 30 minutes, the PBMC layer was collected, washed twice with 1×PBS, and resuspended in 300μL of RPMI-1640 (Hyclone, SH30809.01) supplemented with 10% FBS (Viva cell, C04001-500) and 900μL red blood cell lysis buffer (Solarbio, R1010). The sample was then placed on ice for 15min, followed by centrifugation at 2000 rpm at 4°C for 3 minutes. After washing with 1×PBS, cells were resuspended in 1×PBS for the mitoSplitter experiment. Cell viability was confirmed using Countess™□ Cell Counting Chamber Slides (Invitrogen, C10283).

A549, H2228, H1975, HCC827, and H838 cells were cultured in RPMI-1640 supplemented with 10% FBS and 1% penicillin-streptomycin (Invitrogen, 15140122) at 37°C and 5% CO_2_. For scRNA-seq samples, the five cell lines were co-cultured with equal cell numbers in two culture plates, one with 0.2 μM ARV-771 (MedChemExpress, HY-100972) treatment for 48 hours and another without drug treatment. For bulk mtRNA samples, cells from the five cell lines were cultured independently without drug treatment. To collect cells, the media was removed, and cells were washed with 1×PBS before detachment from the culture plates with Trypsin (Gibco, 25200072) and incubated at room temperature for 2 minutes. After deadhesion, cells were harvested with complete media, washed three times with 1×PBS, and resuspended in 1×PBS for the mitoSplitter experiment.

### Sequencing library preparation for mitoSplitter

The Cell Mitochondria Isolation Kit (Beyotime, C3601) was used to isolate mitochondria from individual samples for bulk mitochondrial RNA (mtRNA) sequencing, following the manufacturer’s instructions. RNA was then purified from the isolated mitochondria using the Total DNA/RNA Isolation Kit (OMEGA, R6731-01). The library preparation for bulk mtRNA sequencing followed a modified Smart-Seq2 protocol(37, 38). In brief, purified mitochondrial RNA (5 μL) was subject to reverse transcription by adding 2 μL 10 μM oligo-dT (ordered from Sangon, **Supplementary Table 12**), 2 μL 10 mM dNTPs (TransGen Biotech, AD101-11), 4 μL 5×RT buffer, 1 μL Maxima H Minus reverse transcriptase (Thermo Scientific, EP0752), 1 μL 50 μM Template Switch oligo (ordered from Sangon, **Supplementary Table 12**), 1 μL RNase Inhibitor (TransGen Biotech, AI101-02), 2 μL 10 mM GTP (Invitrogen, R0461), and 2 μL 50% PEG-8000 (Beyotime, R0056). The reverse transcription was then incubated at 42°C for 90 minutes followed by 5 cycles of 50°C for 2 minutes and 42°C for 2 minutes, and a final incubation at 70°C for 15 minutes. For PCR, 25 μL 2×HiFi HotStart Readymix (Kapa Biosystems, KK2602), 0.8 μL 50 μM ISPCR oligo (ordered from Sangon, **Supplementary Table 12**), and 4.2 μL Nuclease-free water were added. The PCR reaction was incubated at 98°C for 3 minutes followed by 10 cycles of 98°C for 20 seconds, 67°C for 15 seconds, and 72°C for 6 minutes, and a final incubation at 72°C for 6 minutes. The procedure above was independently performed on cells from different donors or cell lines.

For single-cell RNA sequencing, the chip fabrication, barcode bead preparation, and sequencing library preparation were performed as previously described(19). To prepare pseudo-mixed PBMC samples, cells from 11 donors were subject to scRNA-seq independently, and 10,000-dual-well chips were used. To prepare experimentally multiplexed scRNA-seq data, PBMCs from 5 donors were subject to independent and multiplexed scRNA-seq (cells from different donors were mixed thoroughly with the equal input before loading to the Well-paired-seq chip) simultaneously, and 30,000-dual-well chips were used. To generate a large scRNA-seq dataset from 10 donors, PBMCs from 10 donors were subject to multiplexed scRNA-seq, and 100,000-dual-well chips were used for scRNA-seq.

The amplification products of both bulk and scRNA-seq samples were purified twice using 0.6× VAHTS DNA Clean Beads (Vazyme, N411-02), and cDNA concentrations were measured with a Qubit fluorometer. An appropriate volume of cDNA was used as an input for library construction using the TruePrep DNA Library Prep Kit V2 for Illumina (Vazyme, cat#TD502). The reaction was performed following the instructions of the manufacturer except for the primers. For bulk sample and scRNA-seq sample preparation, primer N501 and primer P5-TSO_Hybrid (**Supplementary Table 12**) were used as the P5 primer, respectively. Additionally, primers Nextera_N7×× (**Supplementary Table 12**) were used as the P7 primer in place of the kit’s provided oligonucleotides. The libraries were purified twice using 0.6× VAHTS DNA Clean Beads, and cDNA concentration was measured again. Finally, the libraries of bulk samples and scRNA-seq samples were sequenced on Illumina Novaseq 6000 and HiSeq X Ten instruments with paired-end 150 base reads, respectively.

### Identification of different cell proliferation rates in NSCLC cell lines under ARV-771 treatment

The A549, H2228, H1975, HCC827, and H838 cell lines were cultured, harvested independently, and diluted to 5 cells/μL using complete culture media. Subsequently, 100 μL of cell suspension for each cell line was added to different wells of 96-well plates. After a 24-hour culture period for cell adhesion, each cell line was exposed to ARV-771 at six different doses (0.0625 μM, 0.125 μM, 0.25 μM, 0.5 μM, 1 μM, and 2 μM) in six replicates. The experiment included a total of 42 independently treated cell populations of each cell line, including a vehicle control. After a 48-hour treatment, the cell viability of each condition was measured using the cell counting kit-8 assay (Apexbio, K1018-5). The IC_50_ values of each cell line were calculated to determine the best ARV-771 concentration to result in different cell proliferation rates in NSCLC cell lines.

### Validation of the effect of combined ARV-771 and etoposide on the proliferation of A549 cells

Pre-experiments were conducted to determine the dose range of etoposide that could effectively inhibit the proliferation rate of A549 cells. Subsequently, A549 cells were exposed to ARV-771 and etoposide (MedChemExpress, HY-13629) at the following combinations of doses, 0 μM, 0.0625 μM, 0.125 μM, 0.25 μM, 0.5 μM, 1 μM, and 2 μM of ARV-771, and 0 μM, 0.01 μM, 0.1 μM, 0.2 μM, 0.4 μM, 0.8 μM, and 1 μM of etoposide. This experiment included a total of 48 independently treated cell populations, including vehicle controls, and each combination had six replicates. Cell viability was measured using the CCK-8 assay, and the combination of 0.5 μM ARV-771 and 0.2 μM etoposide was selected for statistical analysis with controls.

### Data pre-processing

To verify the reliability and robustness of mitoSplitter, we collected data from several publicly available single-cell sequencing datasets.

General single-cell sequencing data included three datasets. Dataset 1 was obtained from Gene Expression Omnibus (GEO) with accession number GSE96583(9) and consisted of 3,177 PBMCs from 4 individuals (detailed in **Supplementary Table 1**). Dataset 2 was obtained from GSE213096 and consisted of 25,078 PBMCs from 3 individuals (detailed in **Supplementary Table 2**). Dataset 3 was obtained from the GSA-Human repository with accession number HRA001748 and consisted of 60,000 liver cells from 10 individuals (detailed in **Supplementary Table 3**).

We obtained scRNA-seq data with exogenous barcodes from a previous study(39) (detailed in **Supplementary Table 7**). Detailly, we downloaded datasets generated by CellPlex (European Nucleotide Archive (ENA) repository with accession number E-MTAB-9964), covering 3 samples of 14,606 cells, LMO (custom) (ENA, E-MTAB-11401), covering 4 cell lines of 11,156 cells, TotalSeq-A (ENA, E-MTAB-11401), covering 4 cell lines of 10,171 cells, and TotalSeq-B (ENA, E-MTAB-9964), covering 3 samples of 16,263 cells.

Four sets of sequencing data, including mitochondrial bulk RNA-seq generated using a modified Smart-Seq2 protocol(37, 38), and single-cell RNA-seq data generated from the Well-paired-seq(19) platform, were generated. The first set covered 19,319 PBMCs from 11 individuals (dataset 4, detailed in **Supplementary Table 4**). The second set consisted of 25,275 PBMCs from 5 donors and included independent mitochondrial bulk RNA-seq and single-cell RNA-seq data (included in dataset 5, detailed in **Supplementary Table 8**), as well as a multiplexed single-cell RNA-seq data with 18,944 cells from these 5 donors (included in dataset 5, detailed in **Supplementary Table 8**). The third set was a multiplexed single-cell RNA-seq dataset covering 17,130 PBMCs from 10 donors (dataset 6, detailed in **Supplementary Table 9**). The fourth set consisted of two multiplexed single-cell RNA-seq experiments covering 5 non-small cell lung cancer cell lines under drug and control treatment, with 2,687 drug-treated cells and 2,065 control cells sequenced (detailed in **Supplementary Table 10**).

### Generation of single-cell and bulk mitochondrial mutation matrices

To generate single-cell and bulk mitochondrial mutation matrices, three steps were undertaken: sequence alignment, remapping, and variant calling. The scRNA-seq data were aligned to the GRCh38 genome using CellRanger v3.1.0 (for 10x Genomic RNA-seq data) or zUMIs v2.9.7e(40) (for Well-Paired-Seq data), during the sequence alignment process, and the bulk mitochondria RNA-seq data were quality-filtered using cutadapt v1.18(41) and aligned to the mitochondrial sequence in the GRCh38 genome using STAR v2.7.3a(42). To improve the quality of alignment and reduce the false positive rate of variant calling, the obtained bam files were remapped to the reference genome using minimap2 v2.24(43) with parameters: -ax splice - t 8 -G50k -k 21 -w 11 --sr -A2 -B8 -O12,32 -E2,1 -r200 -p.5 -N20 -f1000,5000 -n2 -m20 -s40 - g2000 -2K50m --secondary=no. The variant calling was performed using a python mtRNA genotyping pipeline(14). In detail, reads in the remapping single-cell bam file with base quality or alignment quality lower than 20 were filtered, and the allele frequency of each cell was then extracted. For the bulk mitochondria RNA-seq bam file, the threshold value of base quality and alignment quality was set to 10. Finally, all the mutations from all samples or cells were integrated to create a bulk or single-cell mutation matrix.

### Demultiplexing of mixed sequencing data using mitoSplitter

We used scanpy v1.9.1(44) to select high-variable SNPs across bulk samples from a mitochondrial mutation matrix for mitochondrial genotyping. These mitochondrial sites were more donor-specific, so we calculated the correlation between single cells and each bulk sample based on the variants frequency of these sites to evaluate the probability of cells from each donor. A single cell was considered to be from the same donor as the bulk sample whose bulk sample exhibited the highest correlation unless it lacked a positively correlated bulk sample. Cells that lacked mitochondrial genotyping sites were unable to receive donor labels through this pipeline due to the uneven coverage of scRNA-seq data. To infer the donor origin of each unbiased cell, we predicted it based on the mtRNA SNPs in scRNA-seq and the inferred labels. We defined this task as a semi-supervised learning problem and used LPA to propagate labels through cells using mtRNA SNPs in scRNA-seq.

In detail, we classified (X, Y) into labeled data as (x_1_, y_1_) … (x_l_, y_l_) and unlabeled data as (x_l+1_, y_l+1_) … (x_l+u_, y_l+u_), depending on whether the sample source labels were obtained. X was the matrix of mtRNA SNPs, and Y was the matrix of cell sample source label, **Y**_**L**_={y_1_, … y_l_} consisted of inferred labels and **Y**_**U**_={y_l+1_ … y_l+u_} was undetermined. The task was to estimate **Y**_**U**_ from **X** and **Y**_**L**_. LPA created a fully connected graph according to the similarity matrix constructed from **X**, where the nodes represented all cells. The edges between nodes were weighted by calculating the normalized graph Laplacian matrix. Then, labeled nodes acted as sources to spread labels **Y**_**L**_ among cells through the edges. Larger edge weights meant easier label propagation. Finally, unlabeled cells received persistent sample labels **Y**_**U**_ when the network of the graph reached a globally stable state. We set the clamping factor α=0.8, which helped the algorithm keep 80% of input labels and change the confidence of labels within 20% during propagation. This optimization was based on inferred labels from unlabeled data. Based on the LPA model, we calculated the probability of each unlabeled cell belonging to each donor. The final cell sample label was determined by the LPA result with a probability greater than (1-10^−10^).

### Data dimension reduction and visualization

The effect of mitochondrial mutation on demultiplexing was illustrated using mtRNA mutation-based TSNE plots. Similar to the workflow for single-cell gene expression, Seurat was used to normalize data, as well as scaling, clustering, and dimension reduction based on highly variable mtRNA SNP features.

### Generation of simulated multiplexed single-cell and bulk RNA-seq data

To simulate multiplexed single-cell RNA-seq data, we utilized a combination of multiple single-cell sequencing datasets. Based on the number of mixed samples *k* and the number of singlets predicted by scrublet v0.2.3(45), we randomly selected the specified number of cells presented within each sample followed by the random selection of 1/*k* of the reads from their corresponding reads for mixing. To remove duplicated reads with the same read ID from different samples, we used the seqkit v2.3.0(46) rmdup function. The remaining reads were mapped to the GRCh38 genome using CellRanger v3.1.0 or zUMIs v2.9.7e(40). After remapping and variant calling, a single-cell mtRNA mutation matrix could be obtained. Finally, the bulk mtRNA mutation matrix was produced by combining the reads of all cells within a sample.

### Computational benchmarking

We compared four demultiplexing algorithms with mitoSplitter: Souporcell v2.0(13), demuxlet v2(9), scSplit v1.0.9(11), and Vireo v0.2.3(12). Each algorithm utilized a maximum of 50 threads, and default pipeline and parameters were used for all demultiplexing calculations. For Souporcell, we ran souporcell_pipeline.py through the singularity container. For demuxlet, we mapped single-cell RNA-seq data from each sample to the GRCh38 genome using CellRanger v3.1.0, generated bam files for genotype calling, and output genotypes through VCF files by samtools v1.9(47) and bcftools v1.8(47). We merged the VCF files into a single file using VCFtools v0.1.17(48), and obtained the demultiplexing results using popscle v0.1-beta demuxlet function, with merged VCF file and “GT” field. For scSplit, we conducted quality-filtering using samtools with “-q 10” and umi_tools v1.1.2(49) dedup function. Variant calling was performed using freebayes v1.2.7(50) with “-iXu -C 2 -q 1” and bcftools filter with “-I ‘%QUAL>30’”. We generated the scSplit count matrix using the scSplit count function and conducted demultiplexing using scSplit run function with “-d 0”.

To conduct performance verification on pooled single-cell sequencing data, we calculated the AUC value, accuracy, and recall of single cells using the Python package sklearn v1.0.2. Singlets predicted by scrublet v0.2.3(45) in each sample were used for performance evaluation. We recorded the running time and memory requirements of the algorithms using custom scripts.

### Cell clustering and annotation

We used Seurat v4.3.0(51) to filter cells with less than 3 features detected or less than 500 UMIs or proportion of mitochondrial genes > 15%, and to perform data normalization, feature selection, scaling, clustering and dimension reduction. Cell type annotation was conducted using celltypist v1.3.0(52) with the “Immune_All_Low.pkl” model. We annotated each Seurat cluster with the top one or two cell types by proportion. For each cell type, we used a paired t-test to compare cell type proportions across samples based on the true and predicted labels. Marker genes for each cluster were identified using the “FindAllMarkers” function in Seurat.

## Data availability

The datasets used in the present study are all publicly available. Dataset 1 was obtained from Gene Expression Omnibus with accession number GSE96583. Dataset 2 was obtained from GSE213096. Dataset 3 was obtained from the GSA-Human repository with accession number HRA001748. Single-cell RNA-seq data with exogenous barcodes were downloaded from European Nucleotide Archive repository, under accession number E-MTAB-9964 for CellPlex and TotalSeq-B, and E-MTAB-11401 for LMO (custom) and TotalSeq-A. Raw Fastq files in dataset 4-6 and non-small cell lung cancer cell lines experiments are available in GSA-human under HRA004390 (https://ngdc.cncb.ac.cn/gsa-human/browse/HRA004390).

## Code availability

The code is available on https://github.com/lnscan/mitoSplitter.

## Acknowledgments

This work was supported by MOST 2020YFA0803601, 2018YFA0801300, NSFC 32071138 and SKLGE-2118 to Jin Li, NSFC 22104080 to Jia Song, and NSFC 21735004, 21927806 to Chaoyong Yang, and Innovative research team of high-level local universities in Shanghai SHSMU-ZLCX20212601.

## Author Contributions

Conceptualization, J.S., C.Y., J.L.; Investigation, L.L., Y.C.; Writing, J.S., X.L., Y.C., L.L., J.L.; Supervision, J.S., J.L., C.Y., X.L., Y.C., L.L., K.Y., R.C., X.W., G.Y., Z.W., Y.Z.

## Conflict of Interest Statements

The authors claim no conflict of interest.

## Notes

### Competing Interest Statement

The authors have declared no competing interest.

### Summary of Updates

Figure 3-5 revised.

