## supplementary tables and figures for "mitoSplitter: A mitochondrial variants-based method for efficient demultiplexing of pooled single-cell RNA-seq"

* Equal contribution

1 Supplementary Note
12 Supplementary Tables
6 Supplementary Figures

### Supplementary Notes

#### Supplementary Note 1: Performance evaluation of demultiplexing tools

Each dataset was validated using the predicted singletons for Demuxlet(1), Souporcell(2), scSplit(3), and Vireo(4). Cells assigned as "Unassigned", "Doublets", or with predicted probability lower than the default threshold were filtered. A predicted label list and a real label list were generated for all remaining cells to evaluate the demultiplexing outcomes. Accuracy was calculated using the 'accuracy_score' function in the Python package sklearn, and recall, precision, and *F1*-score were calculated using the 'recall_score', 'precision_score', and 'f1_score' functions with the parameter 'average = macro'. AUC was computed utilizing the 'preprocess.label', 'metrics.roc_curve', and 'metrics.auc' functions in the Python package sklearn. Furthermore, each tool's running time and memory usage were recorded using custom scripts.

### Supplementary Tables

#### Supplementary Table 1. Number of cells for each sample in virtually multiplexed scRNA-seq dataset 1(1)

| Sample | Number of cells |
|---|---|
| **p1079** | 792 |
| **p1154** | 917 |
| **p1249** | 734 |
| **p1598** | 734 |
| **total** | 3177 |

#### Supplementary Table 2. Number of cells for each sample in virtually multiplexed scRNA-seq dataset 2

| Sample | Number of cells |
|---|---|
| **SRR21523234** | 7859 |
| **SRR21523235** | 9191 |
| **SRR21523236** | 8028 |
| **total** | 25078 |

#### Supplementary Table 3. Number of cells for each sample in virtually multiplexed scRNA-seq dataset 3

| Sample | Number of cells (randomly selected from the original data) |
|---|---|
| **HRR572935** | 6000 |
| **HRR572940** | 6000 |
| **HRR572945** | 6000 |
| **HRR572954** | 6000 |
| **HRR572958** | 6000 |
| **HRR572965** | 6000 |
| **HRR572978** | 6000 |
| **HRR572985** | 6000 |
| **HRR573015** | 6000 |
| **HRR573046** | 6000 |
| **total** | 60000 |

**Supplementary Table 4. Number of cells for 11 PBMC samples (used for pseudo-mixed datasets)**

| Sample | Number of cells |
|---|---|
| **P1** | 1594 |
| **P2 *** | 2295 |
| **P3** | 1399 |
| **P4** | 1142 |
| **P5 *** | 2093 |
| **P6** | 3464 |
| **P7 *** | 1933 |
| **P8 *** | 1700 |
| **P9** | 1331 |
| **P11 *** | 1208 |
| **P12** | 1160 |
| **total** | 19319 |

* Dataset 4 consisted of five samples marked with an asterisk.

**Supplementary Table 5. AUC values of mitoSplitter on pseudo-mixed samples with various sample sizes and combination conditions**

| Sample size | Combination | AUC |
|---|---|---|
| 2 | P11,P1 | 0.9987 |
| | P1,P3 | 1.0000 |
| | P5,P8 | 0.9990 |
| | P11,P4 | 0.9997 |
| | P12,P9 | 0.9989 |
| | P1,P9 | 1.0000 |
| | P2,P6 | 1.0000 |
| | P5,P6 | 0.9978 |
| | P12,P5 | 0.9987 |
| | P2,P6 | 1.0000 |
| 3 | P1,P7,P9 | 0.9990 |
| | P12,P3,P6 | 0.9955 |
| | P1,P2,P6 | 0.9998 |
| | P11,P3,P4 | 0.9989 |
| | P12,P6,P7 | 0.9837 |
| | P1,P3,P5 | 0.9979 |
| | P3,P7,P8 | 0.9993 |
| | P12,P2,P7 | 0.9983 |
| | P2,P3,P7 | 1.0000 |
| | P2,P4,P6 | 1.0000 |
| 4 | P1,P7,P8,P9 | 0.9992 |
| | P2,P3,P5,P7 | 0.9972 |
| | P3,P5,P6,P8 | 0.9970 |
| | P11,P3,P5,P8 | 0.9962 |
| | P1,P2,P3,P7 | 0.9998 |
| | P1,P5,P8,P9 | 0.9985 |
| | P11,P3,P4,P8 | 0.9920 |
| | P12,P1,P3,P9 | 0.9972 |
| | P1,P2,P4,P9 | 1.0000 |
| | P3,P5,P7,P9 | 0.9967 |
| 5 | P1,P3,P5,P6,P7 | 0.9906 |
| | P11,P12,P1,P4,P8 | 0.9924 |
| | P2,P3,P4,P6,P8 | 0.9941 |
| | P11,P12,P2,P4,P7 | 0.9972 |
| | P1,P5,P6,P7,P9 | 0.9913 |
| | P11,P1,P5,P7,P9 | 0.9979 |
| | P1,P6,P7,P8,P9 | 0.9911 |
| | P11,P12,P3,P5,P7 | 0.9931 |
| | P12,P1,P3,P4,P5 | 0.9988 |
| | P11,P2,P4,P5,P9 | 0.9992 |

| | | |
|---|---|---|
| **6** | P11,P12,P2,P4,P5,P6 | 0.9972 |
| | P12,P3,P4,P7,P8,P9 | 0.9930 |
| | P11,P12,P3,P4,P5,P6 | 0.9955 |
| | P12,P1,P2,P5,P6,P8 | 0.9978 |
| | P11,P12,P1,P2,P3,P6 | 0.9961 |
| | P11,P1,P2,P4,P5,P9 | 0.9990 |
| | P12,P1,P2,P3,P5,P8 | 0.9983 |
| | P11,P12,P1,P2,P6,P8 | 0.9969 |
| | P1,P4,P6,P7,P8,P9 | 0.9894 |
| | P11,P12,P3,P6,P7,P9 | 0.9902 |
| **7** | P11,P1,P3,P5,P6,P7,P9 | 0.9924 |
| | P11,P2,P3,P4,P5,P6,P8 | 0.9935 |
| | P11,P1,P2,P3,P6,P8,P9 | 0.9976 |
| | P11,P12,P1,P3,P4,P7,P8 | 0.9927 |
| | P11,P12,P1,P2,P5,P8,P9 | 0.9974 |
| | P11,P12,P4,P5,P6,P7,P9 | 0.9936 |
| | P11,P12,P1,P2,P4,P5,P9 | 0.9977 |
| | P11,P12,P1,P2,P3,P5,P6 | 0.9951 |
| | P11,P12,P1,P3,P4,P7,P9 | 0.9968 |
| | P11,P12,P2,P3,P4,P7,P8 | 0.9929 |
| **8** | P11,P12,P2,P3,P4,P6,P7,P8 | 0.9897 |
| | P11,P12,P1,P4,P6,P7,P8,P9 | 0.9909 |
| | P12,P1,P3,P4,P5,P6,P8,P9 | 0.9946 |
| | P11,P1,P3,P4,P5,P7,P8,P9 | 0.9944 |
| | P1,P2,P3,P4,P5,P6,P7,P8 | 0.9911 |
| | P12,P1,P2,P3,P4,P5,P6,P7 | 0.9947 |
| | P12,P1,P2,P3,P4,P5,P7,P8 | 0.9950 |
| | P12,P1,P2,P4,P5,P7,P8,P9 | 0.9952 |
| | P12,P1,P2,P4,P5,P6,P8,P9 | 0.9948 |
| | P12,P1,P3,P4,P5,P6,P7,P8 | 0.9909 |
| **9** | P12,P3,P2,P4,P5,P6,P7,P8,P9 | 0.9920 |
| | P12,P11,P2,P4,P5,P6,P7,P8,P9 | 0.9915 |
| | P11,P12,P1,P2,P3,P4,P5,P6,P7 | 0.9943 |
| | P11,P12,P1,P2,P3,P5,P6,P8,P9 | 0.9955 |
| | P11,P12,P1,P2,P3,P4,P5,P6,P7 | 0.9943 |
| | P11,P12,P1,P2,P4,P5,P7,P8,P9 | 0.9948 |
| | P11,P1,P2,P3,P4,P5,P6,P7,P9 | 0.9954 |
| | P11,P12,P1,P2,P4,P5,P6,P7,P8 | 0.9914 |
| | P11,P12,P1,P2,P3,P4,P6,P8,P9 | 0.9940 |
| | P11,P12,P1,P3,P4,P5,P7,P8,P9 | 0.9943 |
| **10** | P11,P12,P1,P2,P4,P5,P6,P7,P8,P9 | 0.9924 |
| | P11,P12,P1,P2,P3,P5,P6,P7,P8,P9 | 0.9941 |
| | P11,P12,P1,P2,P4,P3,P6,P7,P8,P9 | 0.9926 |
| | P11,P12,P1,P2,P4,P5,P3,P7,P8,P9 | 0.9948 |

| | | |
|---|---|---|
| | P11,P12,P1,P2,P4,P5,P6,P3,P8,P9 | 0.9946 |
| | P11,P12,P1,P2,P4,P5,P6,P7,P3,P9 | 0.9948 |
| | P11,P12,P1,P2,P4,P5,P6,P7,P8,P3 | 0.9917 |
| | P3,P12,P1,P2,P4,P5,P6,P7,P8,P9 | 0.9927 |
| | P11,P3,P1,P2,P4,P5,P6,P7,P8,P9 | 0.9925 |
| | P11,P12,P3,P2,P4,P5,P6,P7,P8,P9 | 0.9918 |

**Supplementary Table 6. The proportion of cell types in true samples and mitoSplitter-predicted samples in dataset 4**

| Sample | Cell type* | Predicted sample | True sample |
|---|---|---|---|
| **P2** | B cells | 0.1411 | 0.1356 |
| | CMs | 0.1885 | 0.1857 |
| | NCMs | 0.0232 | 0.0220 |
| | NK cells | 0.1606 | 0.1690 |
| | PCs | 0.0167 | 0.0158 |
| | T cells | 0.4698 | 0.4718 |
| **P5** | B cells | 0.3158 | 0.2711 |
| | CMs | 0.0602 | 0.0600 |
| | NCMs | 0.0209 | 0.0182 |
| | NK cells | 0.0526 | 0.0558 |
| | PCs | 0.0167 | 0.0115 |
| | T cells | 0.5338 | 0.5834 |
| **P7** | B cells | 0.1627 | 0.1494 |
| | CMs | 0.1195 | 0.1114 |
| | NCMs | 0.0422 | 0.0364 |
| | NK cells | 0.0763 | 0.0720 |
| | PCs | 0.0040 | 0.0030 |
| | T cells | 0.5954 | 0.6277 |
| **P8** | B cells | 0.1179 | 0.0983 |
| | CMs | 0.1093 | 0.1683 |
| | NCMs | 0.0086 | 0.0058 |
| | NK cells | 0.0872 | 0.0875 |
| | PCs | 0.0270 | 0.0183 |
| | T cells | 0.6499 | 0.6217 |
| **P11** | B cells | 0.0564 | 0.0481 |
| | CMs | 0.3786 | 0.3479 |
| | NCMs | 0.1127 | 0.0993 |
| | NK cells | 0.0737 | 0.0892 |
| | PCs | 0.0058 | 0.0038 |
| | T cells | 0.3728 | 0.4118 |

* CMs, classical monocytes; NCMs, non-classical monocytes; NK cells, natural killer cells; PCs, plasma cells.

**Supplementary Table 7. Number of cells for samples in published experimentally (antibody- or lipid-based) multiplexed scRNA-seq data**(5)

| Antibody- or lipid-based multiplexing methods | Sample | Number of cells |
|---|---|---|
| **CellPlex** | CMO301 | 5264 |
| | CMO302 | 4540 |
| | CMO303 | 4802 |
| | total | 14606 |
| **LMO (custom)** | DU145 | 1284 |
| | MCF7 | 2717 |
| | MDAMB231 | 2274 |
| | PC3 | 4881 |
| | total | 11156 |
| **TotalSeq-A** | DU145 | 1657 |
| | MCF7 | 2983 |
| | MDAMB231 | 2883 |
| | PC3 | 2648 |
| | total | 10171 |
| **TotalSeq-B** | Hashtag8 | 5088 |
| | Hashtag9 | 5815 |
| | Hashtag10 | 5360 |
| | total | 16263 |

**Supplementary Table 8. Number of cells for samples in multiplexed and individually sequenced scRNA-seq data of 5 PBMC samples (in dataset 5)**

| | Sample | Number of cells* |
|---|---|---|
| **Multiplexed** | P408 | 3864 |
| | P409 | 3972 |
| | P410 | 3869 |
| | P411 | 2601 |
| | P412 | 4638 |
| | total | 18944 |
| **Individual** | P408 | 4731 |
| | P409 | 5803 |
| | P410 | 5430 |
| | P411 | 4450 |
| | P412 | 4861 |
| | total | 25275 |

* The number of cells in multiplexed scRNA-seq data was derived from mitoSplitter's prediction results, while the number of cells in individually sequenced scRNA-seq data was derived from zUMIs' kept barcodes output files.

**Supplementary Table 9. Predicted number of cells in 10 demultiplexed samples from dataset 6**

| Sample | Predicted number of cells |
|---|---|
| **P13S** | 1312 |
| **P14S** | 1633 |
| **P15S** | 1747 |
| **P16S** | 1212 |
| **P17S** | 895 |
| **P18S** | 2427 |
| **P19S** | 760 |
| **P20S** | 2306 |
| **P21S** | 973 |
| **P22S** | 1165 |
| **total** | 14430 |

**Supplementary Table 10. Predicted number of cells in the demultiplexed scRNA-seq data of NSCLC cell lines**

| | Sample | Predicted number of cells |
|---|---|---|
| **Control** | A549 | 1031 |
| | H1975 | 374 |
| | H2228 | 265 |
| | H838 | 139 |
| | HCC827 | 256 |
| | total | 2065 |
| **Drug** | A549 | 2425 |
| | H1975 | 55 |
| | H2228 | 63 |
| | H838 | 38 |
| | HCC827 | 106 |
| | total | 2687 |

**Supplementary Table 11. Differentially expressed genes obtained by comparing decreased (subgroup 1) and increased A549 subgroups (subgroup 0,2) after drug treatment**

| Gene_id | p_val | avg_log2FC | pct.1 | pct.2 | p_val_adj |
|---|---|---|---|---|---|
| **TOP2A** | 1.28E-12 | 2.213936954 | 0.501 | 0.544 | 2.57E-09 |
| **CENPF** | 7.99E-06 | 1.942888037 | 0.48 | 0.553 | 0.01598179 |
| **HIST1H1D** | 3.33E-06 | 1.838180527 | 0.399 | 0.459 | 0.00665487 |
| **MKI67** | 5.11E-07 | 1.72396341 | 0.383 | 0.427 | 0.00102184 |
| **TPX2** | 0.0000 | 1.1989 | 0.318 | 0.317 | 1.21E-14 |
| **CENPE** | 0.0000 | 1.0382 | 0.294 | 0.478 | 4.67E-07 |
| **HIST1H1E** | 0.0000 | 0.9557 | 0.311 | 0.376 | 0.01116435 |
| **ANLN** | 0.0000 | 0.9153 | 0.295 | 0.342 | 0.00139351 |
| **KIF23** | 0.0000 | 0.8773 | 0.275 | 0.3 | 1.76E-07 |
| **FP671120.7** | 0.0000 | 0.8526 | 0.993 | 0.993 | 2.42E-75 |
| **CLSPN** | 0.0000 | 0.8514 | 0.341 | 0.507 | 0.0121676 |
| **CCNB2** | 0.0000 | 0.8162 | 0.276 | 0.272 | 1.45E-19 |
| **AURKA** | 0.0000 | 0.8080 | 0.214 | 0.121 | 2.55E-64 |
| **TTK** | 0.0000 | 0.8060 | 0.241 | 0.41 | 1.76E-05 |
| **FAM111B** | 0.0000 | 0.7514 | 0.285 | 0.475 | 2.49E-10 |
| **SGO2** | 0.0000 | 0.7492 | 0.299 | 0.468 | 9.23E-06 |
| **RRM2** | 0.0000 | 0.7451 | 0.289 | 0.47 | 1.30E-05 |
| **BIRC5** | 0.0000 | 0.7046 | 0.332 | 0.516 | 1.89E-07 |
| **CP** | 0.0000 | 0.7006 | 0.71 | 0.341 | 1.40E-50 |
| **KIF4A** | 0.0000 | 0.6353 | 0.245 | 0.46 | 4.81E-16 |
| **CXCL5** | 0.0000 | 0.6231 | 0.894 | 0.616 | 2.62E-47 |
| **MCM4** | 0.0000 | 0.6206 | 0.323 | 0.489 | 0.03829348 |
| **KIF11** | 0.0000 | 0.6159 | 0.287 | 0.473 | 2.11E-07 |
| **HELLS** | 0.0000 | 0.6147 | 0.384 | 0.584 | 1.17E-06 |
| **PTTG1** | 0.0000 | 0.6015 | 0.352 | 0.568 | 4.89E-14 |
| **CKS2** | 0.0000 | 0.5939 | 0.273 | 0.446 | 0.00398011 |
| **CENPU** | 0.0000 | 0.5763 | 0.287 | 0.477 | 1.29E-04 |
| **MIS18BP1** | 0.0000 | 0.5630 | 0.284 | 0.504 | 2.84E-14 |
| **CDKN3** | 0.0000 | 0.5595 | 0.294 | 0.508 | 5.50E-18 |
| **SMC2** | 0.0000 | 0.5464 | 0.281 | 0.322 | 1.13E-05 |
| **DST** | 0.0000 | 0.5457 | 0.937 | 0.633 | 3.61E-48 |
| **ATAD5** | 0.0000 | 0.5448 | 0.3 | 0.468 | 0.00474538 |
| **KNL1** | 0.0000 | 0.5444 | 0.259 | 0.454 | 1.04E-13 |
| **CA12** | 0.0000 | 0.5439 | 0.932 | 0.689 | 2.49E-49 |
| **GOLGB1** | 6.54E-63 | 0.543632795 | 0.923 | 0.597 | 1.31E-59 |
| **PCLAF** | 6.73E-18 | 0.540267879 | 0.355 | 0.576 | 1.35E-14 |
| **TK1** | 2.15E-15 | 0.529831664 | 0.316 | 0.533 | 4.30E-12 |
| **SPDL1** | 7.04E-08 | 0.526628741 | 0.329 | 0.519 | 0.00014072 |
| **ECT2** | 2.63E-09 | 0.52078967 | 0.273 | 0.318 | 5.26E-06 |
| **BLM** | 2.22E-10 | 0.513566486 | 0.255 | 0.442 | 4.45E-07 |

| | | | | | |
|---|---|---|---|---|---|
| **DNAJC9** | 4.01E-10 | 0.509271127 | 0.304 | 0.489 | 8.01E-07 |
| **LCN2** | 2.06E-48 | 0.508147803 | 0.822 | 0.519 | 4.13E-45 |
| **DSN1** | 1.72E-05 | 0.507633767 | 0.235 | 0.396 | 0.03440122 |
| **SOD2** | 1.35E-51 | 0.499276254 | 0.737 | 0.42 | 2.70E-48 |
| **PRC1** | 8.92E-13 | 0.494412729 | 0.228 | 0.419 | 1.78E-09 |
| **SLFN5** | 3.07E-79 | 0.491841236 | 0.781 | 0.418 | 6.14E-76 |
| **AL033397.1** | 6.30E-14 | 0.489306622 | 0.391 | 0.148 | 1.26E-10 |
| **PBK** | 6.39E-20 | 0.485854791 | 0.176 | 0.352 | 1.28E-16 |
| **SOX4** | 5.97E-85 | 0.482556796 | 0.682 | 0.328 | 1.19E-81 |
| **ALCAM** | 1.07E-59 | 0.478943933 | 0.852 | 0.511 | 2.13E-56 |
| **ARL6IP1** | 3.01E-14 | 0.47069325 | 0.364 | 0.579 | 6.03E-11 |
| **MTUS1** | 6.34E-41 | 0.468196869 | 0.713 | 0.394 | 1.27E-37 |
| **CDC20** | 3.00E-12 | 0.467670095 | 0.213 | 0.37 | 5.99E-09 |
| **HIST1H1C** | 4.35E-19 | 0.449064731 | 0.458 | 0.706 | 8.70E-16 |
| **BRCA2** | 8.02E-12 | 0.447835299 | 0.197 | 0.385 | 1.60E-08 |
| **SLC7A11** | 2.00E-47 | 0.446510782 | 0.875 | 0.57 | 4.00E-44 |
| **EZH2** | 2.73E-16 | 0.443538598 | 0.296 | 0.511 | 5.46E-13 |
| **NASP** | 8.97E-07 | 0.442384054 | 0.458 | 0.661 | 0.00179386 |
| **DNMT1** | 6.66E-17 | 0.441894618 | 0.345 | 0.583 | 1.33E-13 |
| **BUB1** | 5.05E-16 | 0.435636594 | 0.252 | 0.456 | 1.01E-12 |
| **RARRES1** | 7.88E-10 | 0.42922574 | 0.372 | 0.12 | 1.58E-06 |
| **CIP2A** | 3.77E-22 | 0.427951835 | 0.217 | 0.431 | 7.54E-19 |
| **CENPN** | 1.09E-18 | 0.424913202 | 0.232 | 0.252 | 2.19E-15 |
| **MAP1B** | 1.36E-38 | 0.424863557 | 0.989 | 0.858 | 2.71E-35 |
| **WDHD1** | 2.54E-18 | 0.421750491 | 0.242 | 0.258 | 5.08E-15 |
| **CENPK** | 1.17E-12 | 0.419986254 | 0.241 | 0.427 | 2.34E-09 |
| **ZKSCAN1** | 1.78E-61 | 0.418933282 | 0.683 | 0.348 | 3.55E-58 |
| **ATP1B1** | 2.92E-33 | 0.416260596 | 0.904 | 0.66 | 5.84E-30 |
| **FAM107B** | 1.04E-44 | 0.411640515 | 0.847 | 0.495 | 2.08E-41 |
| **DCBLD2** | 2.57E-35 | 0.411244266 | 0.957 | 0.697 | 5.14E-32 |
| **TRAM1** | 5.47E-49 | 0.410402426 | 0.858 | 0.542 | 1.09E-45 |
| **CDK1** | 3.76E-09 | 0.405632798 | 0.187 | 0.189 | 7.52E-06 |
| **ARHGAP11A** | 1.90E-28 | 0.405342969 | 0.248 | 0.472 | 3.79E-25 |
| **MIA3** | 2.26E-59 | 0.403946097 | 0.729 | 0.38 | 4.52E-56 |
| **MALAT1** | 2.33E-60 | 0.402243104 | 1 | 0.999 | 4.67E-57 |
| **CCND1** | 8.63E-69 | 0.396132188 | 0.783 | 0.429 | 1.73E-65 |
| **CTNNAL1** | 2.43E-12 | 0.394557833 | 0.371 | 0.581 | 4.86E-09 |
| **RND3** | 1.13E-95 | 0.394473545 | 0.697 | 0.34 | 2.26E-92 |
| **SPTBN1** | 4.48E-62 | 0.387124127 | 0.823 | 0.461 | 8.95E-59 |
| **CDCA8** | 4.51E-09 | 0.37800792 | 0.158 | 0.287 | 9.01E-06 |
| **CTSD** | 3.25E-38 | 0.374216064 | 0.588 | 0.302 | 6.50E-35 |
| **FOS** | 9.22E-69 | 0.372963828 | 0.743 | 0.394 | 1.84E-65 |
| **SGO1** | 2.77E-17 | 0.370450005 | 0.206 | 0.383 | 5.55E-14 |
| **LBH** | 1.04E-80 | 0.368670828 | 0.75 | 0.399 | 2.08E-77 |

| | | | | | |
|---|---|---|---|---|---|
| **AURKB** | 8.12E-09 | 0.364760089 | 0.106 | 0.024 | 1.62E-05 |
| **CKAP2** | 1.03E-17 | 0.363164744 | 0.268 | 0.477 | 2.06E-14 |
| **FGA** | 6.34E-13 | 0.362969017 | 0.5 | 0.266 | 1.27E-09 |
| **CCPG1** | 1.38E-31 | 0.362325201 | 0.726 | 0.427 | 2.76E-28 |
| **SPP1** | 5.49E-39 | 0.361238836 | 0.753 | 0.443 | 1.10E-35 |
| **SAT1** | 7.24E-25 | 0.360370202 | 0.978 | 0.835 | 1.45E-21 |
| **NRCAM** | 1.61E-52 | 0.360254084 | 0.723 | 0.394 | 3.23E-49 |
| **VRK1** | 1.27E-13 | 0.359739244 | 0.235 | 0.427 | 2.53E-10 |
| **CYP1B1** | 3.93E-67 | 0.352294193 | 0.655 | 0.321 | 7.87E-64 |
| **FEN1** | 5.84E-10 | 0.351640077 | 0.187 | 0.366 | 1.17E-06 |
| **CNTN1** | 1.76E-48 | 0.348384584 | 0.423 | 0.119 | 3.52E-45 |
| **TGM2** | 2.72E-32 | 0.346110945 | 0.589 | 0.31 | 5.45E-29 |
| **PTMA** | 3.33E-17 | 0.343902878 | 0.898 | 0.982 | 6.67E-14 |
| **LIMCH1** | 6.34E-41 | 0.341271091 | 0.826 | 0.519 | 1.27E-37 |
| **TEAD1** | 2.14E-55 | 0.33974007 | 0.782 | 0.44 | 4.27E-52 |
| **CCDC34** | 1.40E-11 | 0.336983157 | 0.177 | 0.223 | 2.80E-08 |
| **CTSB** | 6.61E-49 | 0.334988893 | 0.806 | 0.475 | 1.32E-45 |
| **MAP7** | 1.28E-36 | 0.334861406 | 0.42 | 0.139 | 2.56E-33 |
| **AKR1B1** | 1.34E-17 | 0.334842108 | 0.951 | 0.779 | 2.67E-14 |
| **SMC1A** | 4.77E-07 | 0.333385883 | 0.339 | 0.521 | 0.00095374 |
| **GINS2** | 9.33E-09 | 0.332659153 | 0.166 | 0.339 | 1.87E-05 |
| **ANP32E** | 1.33E-12 | 0.331502887 | 0.288 | 0.322 | 2.65E-09 |
| **SKAP2** | 6.43E-72 | 0.331395168 | 0.649 | 0.287 | 1.29E-68 |
| **BICC1** | 6.09E-48 | 0.329204449 | 0.665 | 0.345 | 1.22E-44 |
| **CDC6** | 4.57E-35 | 0.324190045 | 0.205 | 0.437 | 9.14E-32 |
| **CKAP5** | 1.98E-23 | 0.320700757 | 0.334 | 0.561 | 3.96E-20 |
| **TFPI** | 1.52E-09 | 0.320662 | 0.492 | 0.269 | 3.05E-06 |
| **THSD7A** | 3.17E-27 | 0.316512031 | 0.501 | 0.248 | 6.33E-24 |
| **ZNF292** | 1.21E-81 | 0.316032489 | 0.585 | 0.261 | 2.42E-78 |
| **LPP** | 4.62E-11 | 0.315942532 | 0.502 | 0.282 | 9.24E-08 |
| **UTRN** | 4.11E-78 | 0.315598855 | 0.611 | 0.277 | 8.23E-75 |
| **PHLDB2** | 4.03E-69 | 0.315548435 | 0.663 | 0.332 | 8.06E-66 |
| **DTL** | 1.97E-32 | 0.315289015 | 0.141 | 0.347 | 3.94E-29 |
| **NEK2** | 3.81E-34 | 0.31503783 | 0.125 | 0.315 | 7.62E-31 |
| **MCM10** | 2.91E-18 | 0.314071986 | 0.237 | 0.42 | 5.81E-15 |
| **AR** | 1.15E-83 | 0.313996153 | 0.646 | 0.31 | 2.29E-80 |
| **ANKRD30A** | 1.04E-39 | 0.313946829 | 0.443 | 0.168 | 2.07E-36 |
| **TSPAN14** | 5.50E-11 | 0.313355792 | 0.464 | 0.256 | 1.10E-07 |
| **GOLGA4** | 7.56E-25 | 0.313000834 | 0.887 | 0.64 | 1.51E-21 |
| **KIF15** | 4.45E-23 | 0.30684297 | 0.135 | 0.298 | 8.91E-20 |
| **LIMA1** | 9.23E-20 | 0.306533759 | 0.412 | 0.177 | 1.85E-16 |
| **USP1** | 6.33E-17 | 0.305997554 | 0.226 | 0.443 | 1.27E-13 |
| **GEN1** | 1.01E-10 | 0.302081496 | 0.229 | 0.414 | 2.02E-07 |
| **CLDN1** | 3.89E-75 | 0.30163662 | 0.719 | 0.383 | 7.77E-72 |

| | | | | | |
|---|---|---|---|---|---|
| **ALDH1A1** | 2.84E-26 | 0.299234891 | 0.994 | 0.918 | 5.68E-23 |
| **CST1** | 2.58E-06 | 0.298551856 | 0.302 | 0.123 | 0.00516782 |
| **ZC3HAV1** | 4.46E-81 | 0.297749933 | 0.558 | 0.216 | 8.92E-78 |
| **POLQ** | 1.50E-12 | 0.29578892 | 0.216 | 0.375 | 3.00E-09 |
| **XRCC2** | 7.64E-23 | 0.29494689 | 0.192 | 0.406 | 1.53E-19 |
| **AKAP9** | 3.26E-26 | 0.289367512 | 0.754 | 0.478 | 6.51E-23 |
| **UBE2T** | 6.59E-22 | 0.288784087 | 0.225 | 0.447 | 1.32E-18 |
| **CXCL8** | 3.99E-55 | 0.288016257 | 0.761 | 0.427 | 7.98E-52 |
| **PDE4D** | 9.80E-40 | 0.286967288 | 0.545 | 0.232 | 1.96E-36 |
| **JUN** | 3.55E-37 | 0.286569819 | 0.876 | 0.568 | 7.10E-34 |
| **TPM1** | 1.15E-26 | 0.286420871 | 0.686 | 0.408 | 2.31E-23 |
| **MET** | 2.69E-61 | 0.278162076 | 0.602 | 0.282 | 5.37E-58 |
| **ANXA4** | 7.39E-12 | 0.277948251 | 0.611 | 0.385 | 1.48E-08 |
| **DAB2** | 6.95E-32 | 0.277435871 | 0.5 | 0.229 | 1.39E-28 |
| **C1S** | 7.52E-06 | 0.275686143 | 0.396 | 0.189 | 0.0150448 |
| **CCN2** | 4.72E-90 | 0.275664205 | 0.55 | 0.237 | 9.43E-87 |
| **MELK** | 7.78E-07 | 0.274100896 | 0.218 | 0.351 | 0.0015551 |
| **MT2A** | 3.63E-10 | 0.274087408 | 0.238 | 0.433 | 7.27E-07 |
| **TAF9** | 2.32E-25 | 0.273074661 | 0.332 | 0.572 | 4.65E-22 |
| **TIMELESS** | 2.95E-28 | 0.266765494 | 0.23 | 0.465 | 5.90E-25 |
| **SYT1** | 2.82E-51 | 0.264758632 | 0.517 | 0.23 | 5.63E-48 |
| **BRIP1** | 4.60E-25 | 0.26399588 | 0.132 | 0.336 | 9.20E-22 |
| **TNFAIP2** | 4.17E-29 | 0.261893143 | 0.927 | 0.685 | 8.35E-26 |
| **MYO6** | 6.59E-23 | 0.261502545 | 0.471 | 0.226 | 1.32E-19 |
| **ASPH** | 9.49E-20 | 0.260868051 | 0.981 | 0.826 | 1.90E-16 |
| **SLF1** | 1.89E-19 | 0.259977715 | 0.21 | 0.433 | 3.78E-16 |
| **CLDN2** | 1.04E-21 | 0.258339828 | 0.12 | 0.048 | 2.07E-18 |
| **GCNT3** | 1.06E-05 | 0.258129214 | 0.383 | 0.153 | 0.02123725 |
| **FGG** | 4.39E-19 | 0.255840946 | 0.687 | 0.432 | 8.78E-16 |
| **TOP2B** | 1.61E-56 | 0.254302557 | 0.612 | 0.296 | 3.22E-53 |
| **KRT8** | 1.11E-19 | 0.253408662 | 0.997 | 0.938 | 2.23E-16 |
| **HIST1H2AI** | 7.74E-43 | 0.253332286 | 0.132 | 0.372 | 1.55E-39 |
| **DAAM1** | 1.91E-74 | 0.252366802 | 0.663 | 0.329 | 3.82E-71 |

**Supplementary Table 12. Oligonucleotide sequences used in this study**

| Oligonucleotide | Sequence |
|---|---|
| **oligo-dT** | AAGCAGTGGTATCAACGCAGAGTACT$_{30}$VN |
| **Template Switch oligo** | AAGCAGTGGTATCAACGCAGAGTACAT /rG/rG/iXNA_G |
| **ISPCR oligo** | AAGCAGTGGTATCAACGCAGAGT |
| **N501** | AATGATACGGCGACCACCGAGATCTACACTAGATCGCTCGTCGGCAGCGTC |
| **P5-TSO_Hybrid** | AATGATACGGCGACCACCGAGATCTACACGCCTGTCCGCGGAAGCAGTGGTATCAACGCAGAGT-s-A-s-C |
| **Nextera_N701** | CAAGCAGAAGACGGCATACGAGATTCGCCTTAGTCTCGTGGGCTCGG |
| **Nextera_N702** | CAAGCAGAAGACGGCATACGAGATCTAGTACGGTCTCGTGGGCTCGG |
| **Nextera_N703** | CAAGCAGAAGACGGCATACGAGATTTCTGCCTGTCTCGTGGGCTCGG |
| **Nextera_N704** | CAAGCAGAAGACGGCATACGAGATGCTCAGGAGTCTCGTGGGCTCGG |
| **Nextera_N705** | CAAGCAGAAGACGGCATACGAGATAGGAGTCCGTCTCGTGGGCTCGG |
| **Nextera_N706** | CAAGCAGAAGACGGCATACGAGATCATGCCTAGTCTCGTGGGCTCGG |
| **Nextera_N707** | CAAGCAGAAGACGGCATACGAGATGTAGAGAGGTCTCGTGGGCTCGG |
| **Nextera_N708** | CAAGCAGAAGACGGCATACGAGATCCTCTCTGGTCTCGTGGGCTCGG |
| **Nextera_N709** | CAAGCAGAAGACGGCATACGAGATAGCGTAGCGTCTCGTGGGCTCGG |
| **Nextera_N710** | CAAGCAGAAGACGGCATACGAGATCAGCCTCGGTCTCGTGGGCTCGG |

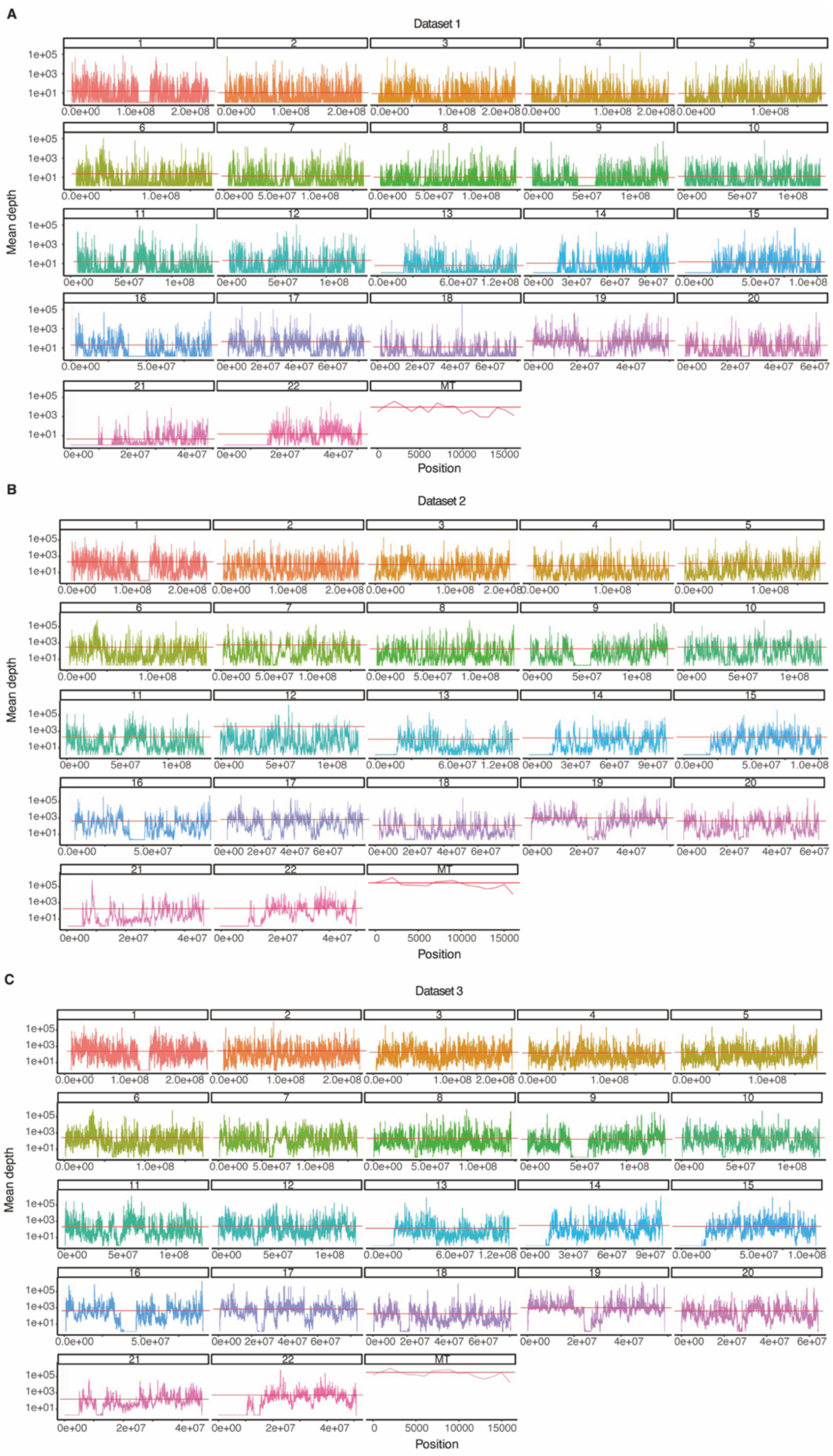


**Supplementary Fig. 1 | Comparison of the sequencing depth of the mitochondrial genome to that of the autosomes.** (A) Dataset 1 contains 3,177 cells from 4 samples(1). (B) Dataset 2 contains 25,078 cells from 3 samples. (C) Dataset 3 contains 60,000 cells from 10 samples. The mean sequencing depth of all sites within a 1 kbp slide window was calculated.

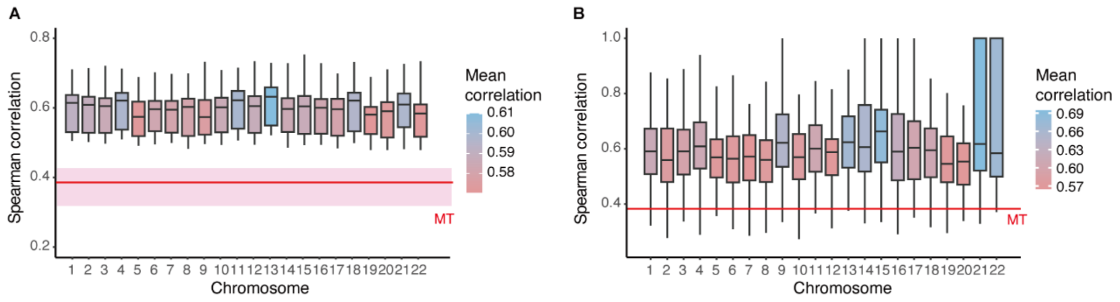


**Supplementary Fig. 2 | Mitochondrial variants are the optimal material for genotyping.** (A) The Mean Spearman correlation between 5000 randomly selected pairs of two individuals in 1000 genomes project (phase 3)(6) based on mitochondrial or autosomal variant profiles. The red line represents the mean correlation of the mitochondrial variants between individuals, the upper and lower pink boundaries represent the first and third quantiles. MT represents “mitochondrion”. It revealed that mitochondrial variants profile is significantly more variable between individuals than that of other chromosomes. (B) Mean Spearman correlation between pairwise individuals from 1000 genomes project (phase 3) based on variant profiles of mitochondrial or autosomal segments. The length of an autosomal segment is equal to the length of the mitochondrial genome, and 100 segments were randomly extracted from each autosomal. The red line represents the mean correlation of the mitochondrial genome between individuals. MT represents “mitochondrion”. It revealed that mitochondrial variant profiles are significantly more variable than those of other chromosomes within the same length interval (1 mitochondrial length).

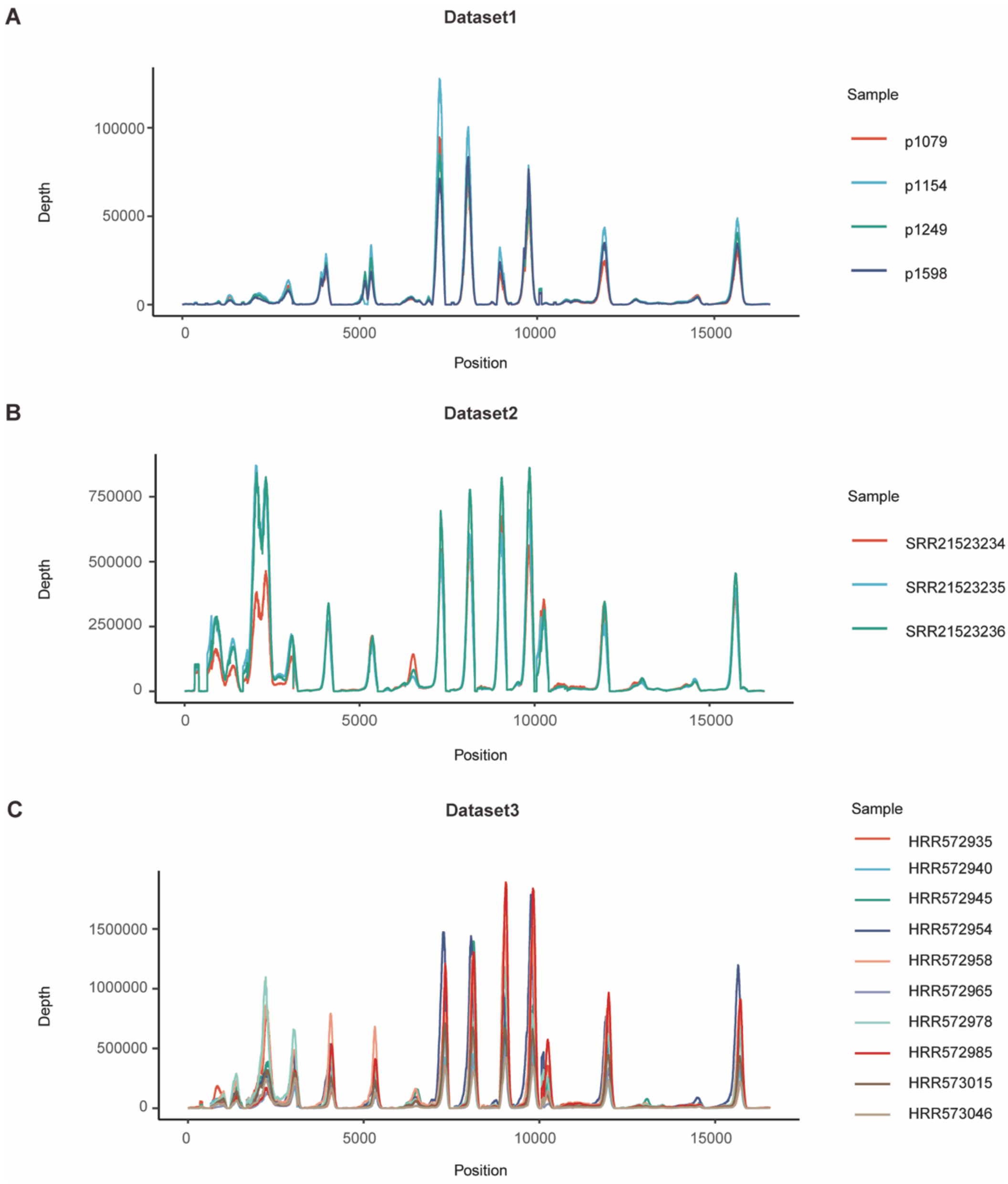


**Supplementary Fig. 3 | Sequencing depth of scRNA-seq data in the mitochondrial genome for datasets 1-3.** (A) Dataset 1 contains 3,177 cells from 4 samples(1). (B) Dataset 2 contains 25,078 cells from 3 samples. (C) Dataset 3 contains 60,000 cells from 10 samples. Each sample was represented by a line that displayed the total sequencing depth for each base of the mitochondrial genome.

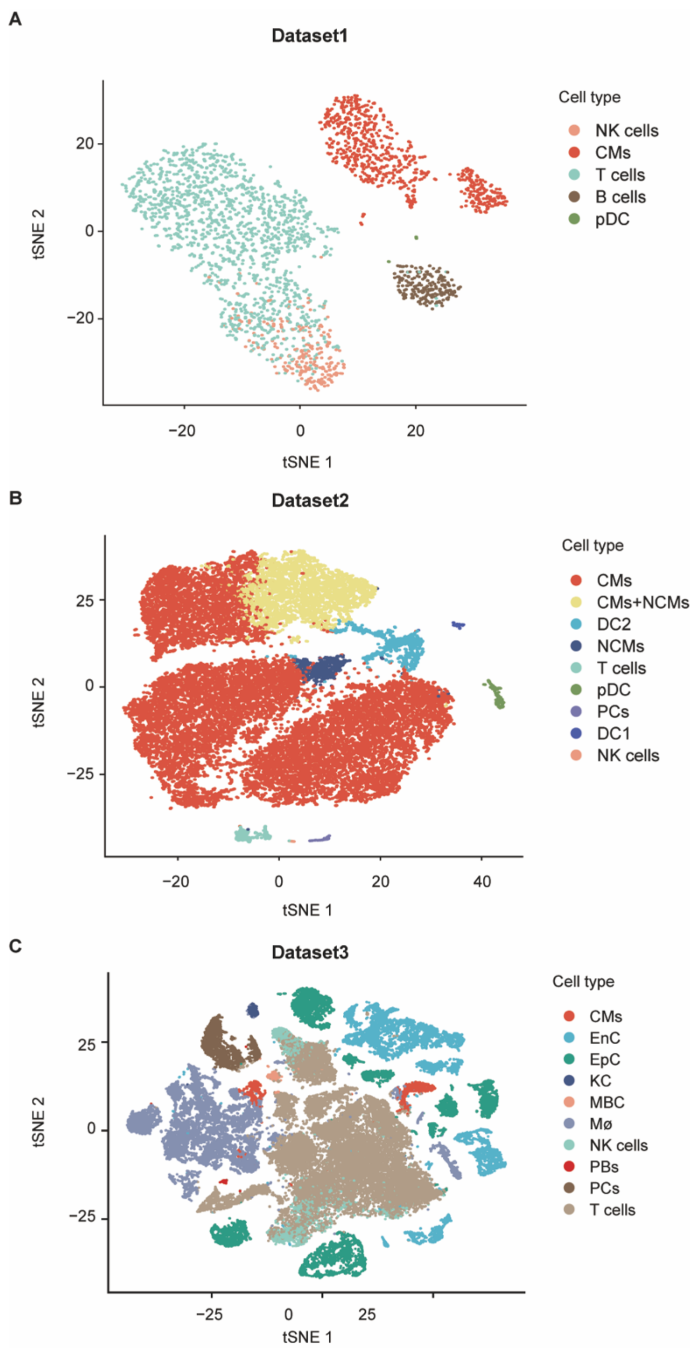


**Supplementary Fig. 4 | The annotated cell types in the virtually multiplexed scRNA-seq data.** (A) Dataset 1 contains 3,177 cells from 4 samples(1). (B) Dataset 2 contains 25,078 cells from 3 samples. (C) Dataset 3 contains 60,000 cells from 10 samples. The cell types were annotated using celltypist v1.3.0(7). CMs, classical monocytes; NCMs, non-classical monocytes; PCs, plasma cells; MBC, memory B cells; NK cells, natural killer cells; pDC, plasmacytoid dendritic cells; DC1, type 1 dendritic cells; DC2, type 2 dendritic cells; PBs, plasmablasts; Mø, Macrophages; EnC, Endothelial cells; EpC, Epithelial cells; KC, Kupffer cells.

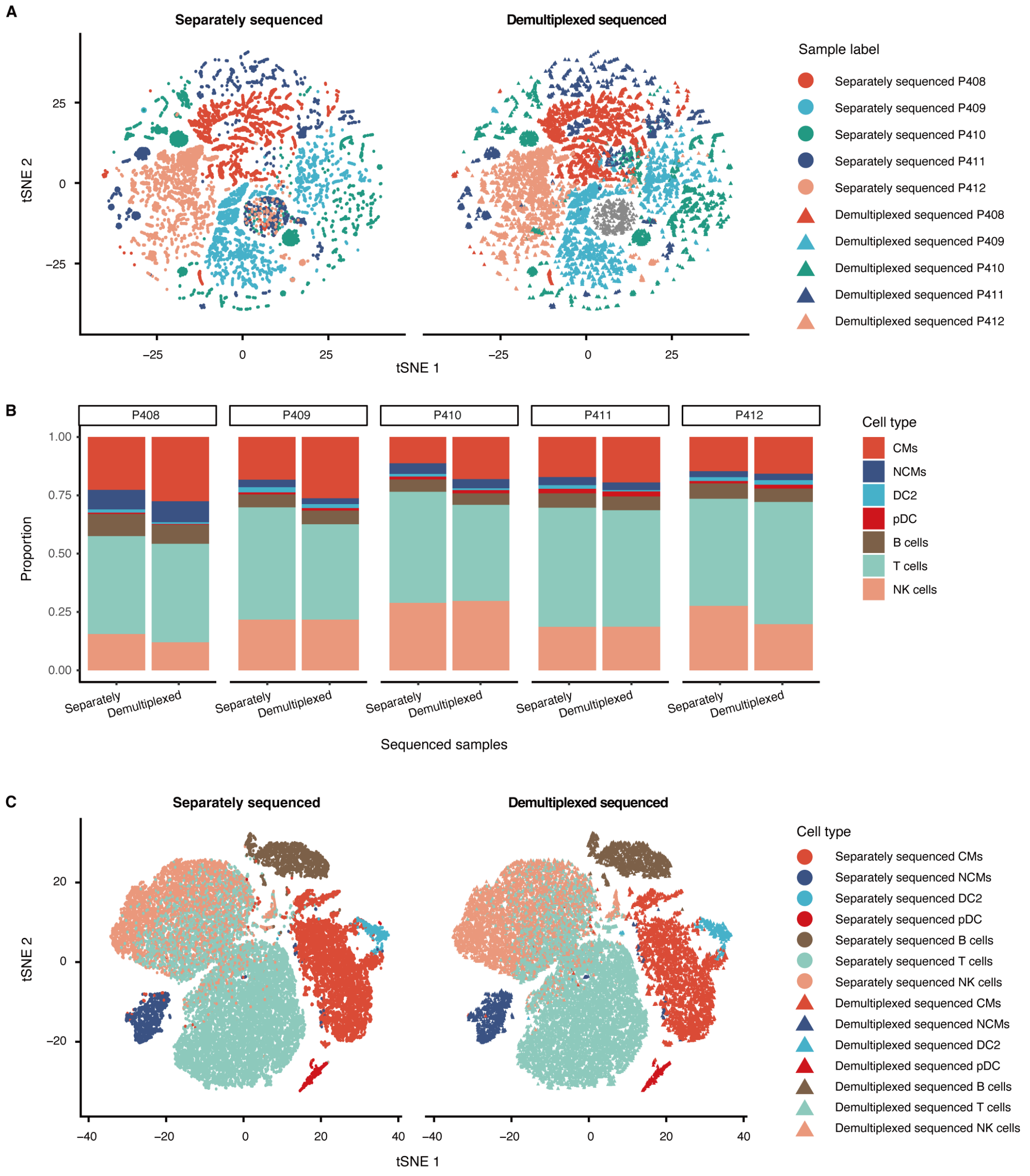


**Supplementary Fig. 5 | Performance of mitoSplitter on experimentally multiplexed dataset 5.** (A) TSNE plot derived from highly variable variants identified by mitoSplitter and colored according to sample labels, the light circles represent separately sequenced sample cells, and the dark triangles represent demultiplexed cells. (B) The proportion of each cell type in the separately sequenced and demultiplexed samples. (C) Gene expression-based TSNE plot for separately sequenced and demultiplexed data colored in cell types annotated using celltypist v1.3.0(7). CMs, classical monocytes; NCMs, non-classical monocytes; DC2, type 2 dendritic cells; pDC, plasmacytoid dendritic cells; NK cells, natural killer cells.

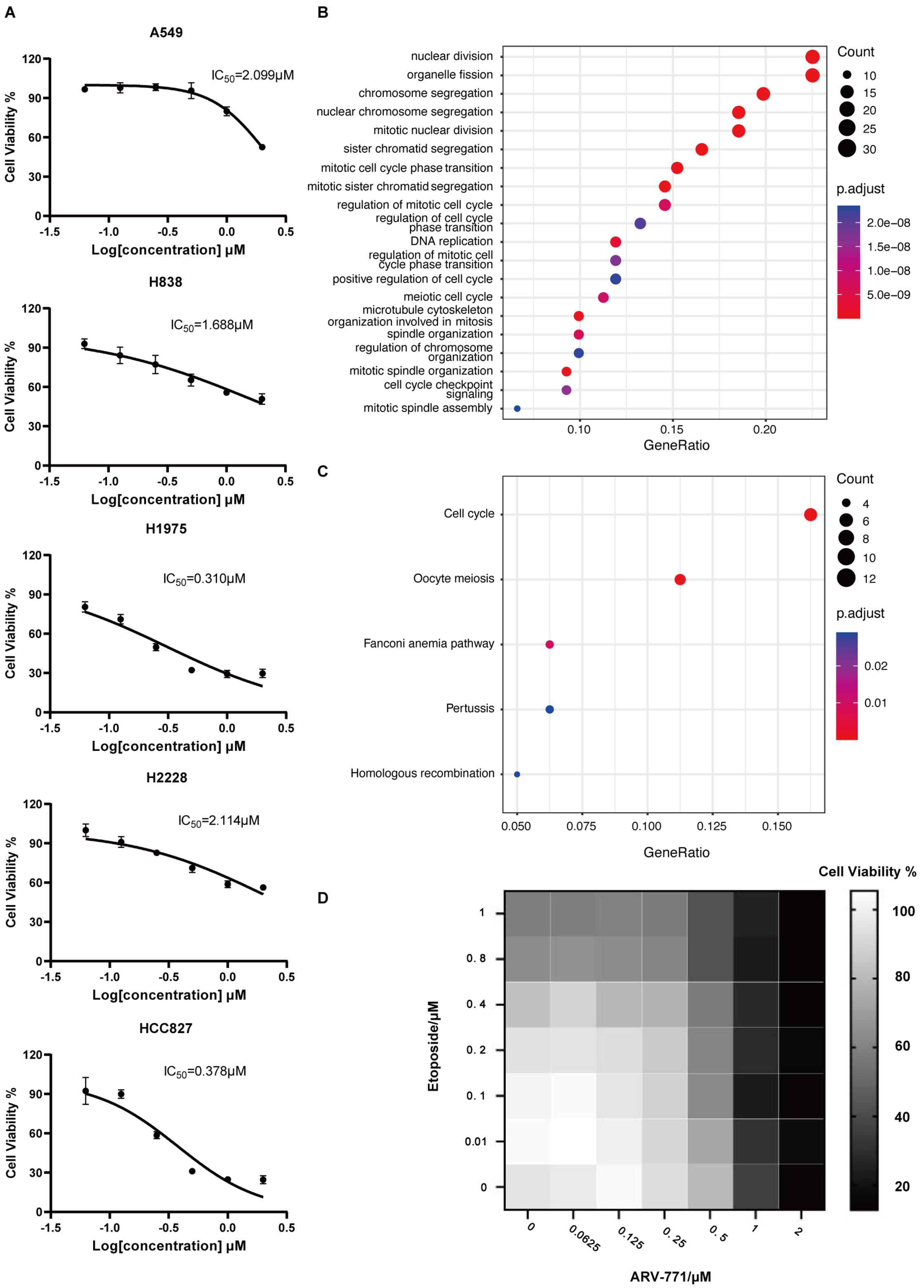


**Supplementary Fig. 6 | Multiplexed single-cell RNA-seq analysis of NSCLC by mitoSplitter.** (A) The chemical degradation of the BET via the treatment of ARV-771 had different effects on the proliferation rate of NSCLC cell lines. (B) Top 20 enriched terms of biological processes in Gene Ontology (GO) enrichment analysis. (C) Enriched pathways in KEGG enrichment analysis. (D) The effect of combined administration of two drugs at different concentrations on cell viability.
